# PRISMA: A tensor-based framework for deconstructing the genetic architecture of complex diseases, with application to diabetic retinopathy

**DOI:** 10.64898/2026.05.25.727382

**Authors:** Hao Xiong, Wenbin Xu, Aocheng Ji, Li Zhong, Siyan Liu, Ziwei Xie, Jing Yan, Zhengzheng Wu

## Abstract

Complex-disease GWAS compress tissue-dependent genetic effects into aggregate locus-level statistics, obscuring regulatory trajectories that shape heterogeneous disease biology. We developed PRISMA (Polygenic Risk Integration via Summary-statistics Multi-tissue Array-decomposition), an LD-aware summary-statistics framework that integrates GWAS and multi-tissue cis-eQTL evidence through graph Laplacian-regularized block-wise factorization. Applied to diabetic retinopathy, PRISMA deconvolved polygenic risk into three tissue-biased axes reflecting vascular-metabolic, systemic immune-inflammatory, and retina-specific neurodegenerative trajectories. The framework prioritized 549 axis-associated targets, including 403 below conventional genome-wide significance, and produced sharper tissue-regulatory resolution than PCA, NMF, and K-means. MAGMA, SMR/HEIDI, and targeted colocalization supported gene-level concordance with established post-GWAS evidence, while a height GWAS control supported trait-dependent regulatory reprioritization. Independent single-cell atlases mapped the axes to fibrovascular, immune, and retinal compartments, and exploratory vitreous proteomic and metabolomic profiling nominated downstream molecular correlates. PRISMA provides a scalable, open-source framework for reframing GWAS interpretation from aggregate locus discovery toward tissue-resolved genetic trajectory mapping.

## Main

Complex diseases, such as diabetic retinopathy (DR), exhibit substantial inter-patient heterogeneity in clinical presentation, progression, and treatment response. This phenotypic diversity likely reflects distinct underlying regulatory mechanisms—a convergent architecture wherein multiple divergent genetic pathways ultimately funnel into a common disease endpoint, consistent with the omnigenic model proposed by Boyle et al. (2017) [1]. However, traditional genome-wide association studies (GWAS) aggregate genetic effects across all patients, providing only averaged signals that obscure disease subtypes and limit mechanistic interpretation and genetically informed stratification hypotheses.

Over the past decade, integrative frameworks have progressively refined our understanding of the genetic architecture of complex diseases. Early methods like summary-data-based Mendelian randomization (SMR) and transcriptome-wide association studies (TWAS) successfully shifted the paradigm from single-variant associations to gene-level mapping by integrating GWAS with expression quantitative trait loci (eQTL) data [2,3]. Building on this foundation, subsequent methods have addressed complementary aspects of genetic architecture. Stratified LD score regression (S-LDSC) partitions disease heritability across regulatory annotations [4], while multivariate fine-mapping methods like mvSuSiE isolate independent causal variants across pleiotropic traits [5]. These approaches excel at their respective objectives—aggregate heritability partitioning and causal variant identification—but are not designed to decompose a single disease into divergent, tissue-anchored genetic risk trajectories at locus-by-tissue resolution.

More directly relevant to trajectory-level decomposition are methods that apply dimensionality reduction to multi-tissue or multi-modal data. MOFA+ performs unsupervised factor analysis to integrate multi-omics data [6], while sn-spMF applies sparse matrix factorization to GTEx eQTL data [7]. While these methods demonstrate the power of latent factor models, they are not specifically designed to incorporate block-wise linkage disequilibrium (LD) topology in summary-level GWAS–eQTL decomposition. Variants within LD blocks share coordinated regulatory effects due to physical co-inheritance, yet standard factorizations treat all variants as independent observations. Without explicit LD topology constraints, the solution space remains underconstrained, potentially yielding axes that lack biological anchoring.

To address this gap, we introduce PRISMA (Polygenic Risk Integration via Summary-statistics Multi-tissue Array-decomposition), a computational framework for decomposing tissue-resolved genetic architecture from GWAS and multi-tissue eQTL summary statistics [8]. PRISMA is formulated as a general locus-by-tissue-by-phenotype framework, but the present study deliberately evaluates its single-phenotype instantiation in diabetic retinopathy. In this setting, the phenotype mode is fixed at *P* = 1, making the empirical model an LD-aware, graph-regularized decomposition of locus-by-tissue genetic architecture rather than a multi-trait loading analysis. This design allowed us to perform a deeper biological and methodological evaluation within one clinically heterogeneous disease while retaining a mathematical specification that can be extended to multi-phenotype applications in future work. PRISMA’s key innovations therefore lie not in tensor dimensionality alone, but in three architectural features: block-wise decomposition that preserves local LD structure across genomic blocks, graph Laplacian regularization anchored to empirical LD topology [9], and non-negativity constraints with warm-up heuristics that support interpretable and empirically stable tissue-biased axes. Unlike aggregate heritability methods that summarize tissue contributions across all loci, or fine-mapping and colocalization methods that evaluate variant-level or local shared-variant evidence without decomposing a disease into regulatory trajectories, PRISMA quantifies how individual loci contribute to tissue-anchored genetic profiles. The resulting axes provide ranked loci and tissue-specificity profiles for downstream biological projection and comparator analysis, while remaining hypothesis-generating rather than causal by themselves.

Here, we applied PRISMA to dissect the genetic architecture of DR, a major cause of vision impairment and blindness worldwide that remains clinically heterogeneous despite advances in screening and treatment [10,11]. PRISMA resolved the DR-associated polygenic signal into three tissue-biased genetic axes: a systemic vascular-metabolic axis, a circulating immune-inflammatory axis, and a retina-specific neurodegenerative axis. We evaluated these axes using complementary evidence layers: conventional dimensionality-reduction baselines, established post-GWAS comparator analyses including MAGMA, SMR/HEIDI, and targeted colocalization, a height GWAS negative-control rerun, independent single-cell transcriptomic atlases, and exploratory vitreous humor proteomic and metabolomic profiling. Together, these analyses position PRISMA as a single-trait proof-of-principle for LD-aware, tissue-resolved decomposition of complex-disease genetic architecture. The inferred axes should be interpreted as genetically anchored regulatory hypotheses and prioritization structures, not as validated causal endotypes or clinical biomarkers.

## Results

### Overview of the PRISMA framework

To overcome the limitations of tissue-collapsed analytical paradigms, we developed PRISMA, a computational framework that preserves the multi-dimensional regulatory landscape of complex traits (Fig. 1). Rather than compressing pleiotropic effects into a single global metric, PRISMA models disease architecture as an interaction between genetic variants, tissue-specific expression profiles, and phenotypic traits. First, PRISMA integrates GWAS summary statistics with multi-tissue eQTL data to derive a standardized association statistic (*S*_*int*_, GWAS-eQTL integration score) for each genetic variant and tissue. These scores are projected into a variant-by-tissue association tensor (𝒳). PRISMA then applies graph-regularized block-wise alternating least squares (ALS) to approximate the observed tensor through low-rank latent factors:

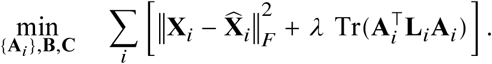

where **A**_𝑖_ (Factor A) represents locus-level genetic drivers (Supplementary Data 1), while global factors **B** (Factor B) and **C** (Factor C) capture tissue-specific regulatory context and phenotypic relevance, respectively (Supplementary Table 12). In the current DR application, the framework is applied to a single phenotype (P=1), so **C** is a rank-specific scaling vector; the P>1 setting remains a future model specification. To preserve local linkage disequilibrium (LD) structure, PRISMA incorporates a graph Laplacian penalty (**L**_𝑖_), encouraging SNPs in high LD to have coordinated factor loadings. Ultimately, each derived component (or “Axis”) represents a distinct genetic risk trajectory linking variant sets to tissue or spatial microenvironments.

**Fig. 1.**
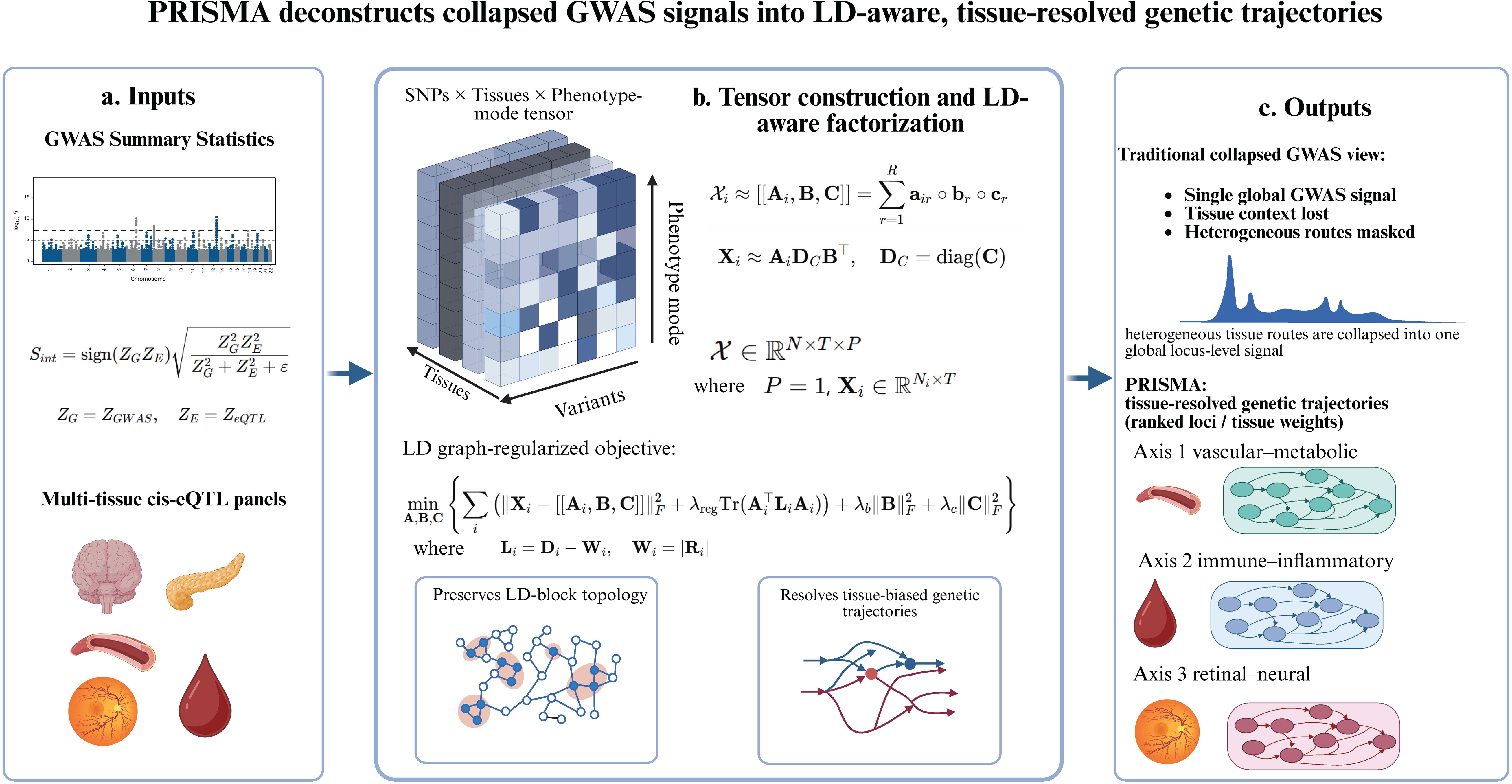
Overview of the PRISMA framework. a, PRISMA integrates GWAS summary statistics with multi-tissue cis-eQTL references to compute a PRISMA-specific GWAS-eQTL integration score (*S*_*int*_), prioritizing variant-gene-tissue entries jointly supported by disease association and local expression regulation. b, Integration scores are organized into a general SNP × tissue × phenotype-mode tensor. In the current diabetic retinopathy application, the phenotype mode is a singleton dimension (*P* = 1), yielding LD-block-specific variant-by-tissue matrices. PRISMA applies graph Laplacian-regularized block-wise factorization to preserve local LD topology while extracting tissue-biased latent axes. c, Unlike conventional GWAS, which collapses heterogeneous tissue routes into one global association signal, PRISMA resolves ranked loci and tissue weights into interpretable genetic trajectories, including vascular-metabolic, immune-inflammatory, and retina-specific neurodegenerative axes. DR, diabetic retinopathy; LD, linkage disequilibrium; eQTL, expression quantitative trait locus; GWAS, genome-wide association study.

This formulation was designed to preserve two features that are often lost in tissue-collapsed GWAS interpretation: the block-wise correlation structure among nearby variants and the tissue context of regulatory effects. Rather than treating each SNP-tissue entry as an independent observation, PRISMA uses the LD graph to regularize local factor loadings while estimating global tissue factors across all blocks. The resulting axes therefore summarize coordinated locus-by-tissue patterns rather than isolated lead-variant associations.

PRISMA showed efficient genome-wide runtime. The block-wise ALS implementation processed 993,226 pre-LD candidate variants across five eQTL tissues (4,966,130 variant-by-tissue cells) with a median wall-clock runtime of 206.3 seconds across five full real-data runs on an Intel Core i9 14900K processor (Supplementary Table 22), supporting practical use on genome-wide summary-statistics data. Monte Carlo simulations supported calibration and recovery under controlled settings, with conservative Type I error calibration and signal-dependent recovery power (78.96% at signal level 2.5 and 95.25% at signal level 3.5; Fig. 2, Supplementary Note 1).

**Fig. 2.**
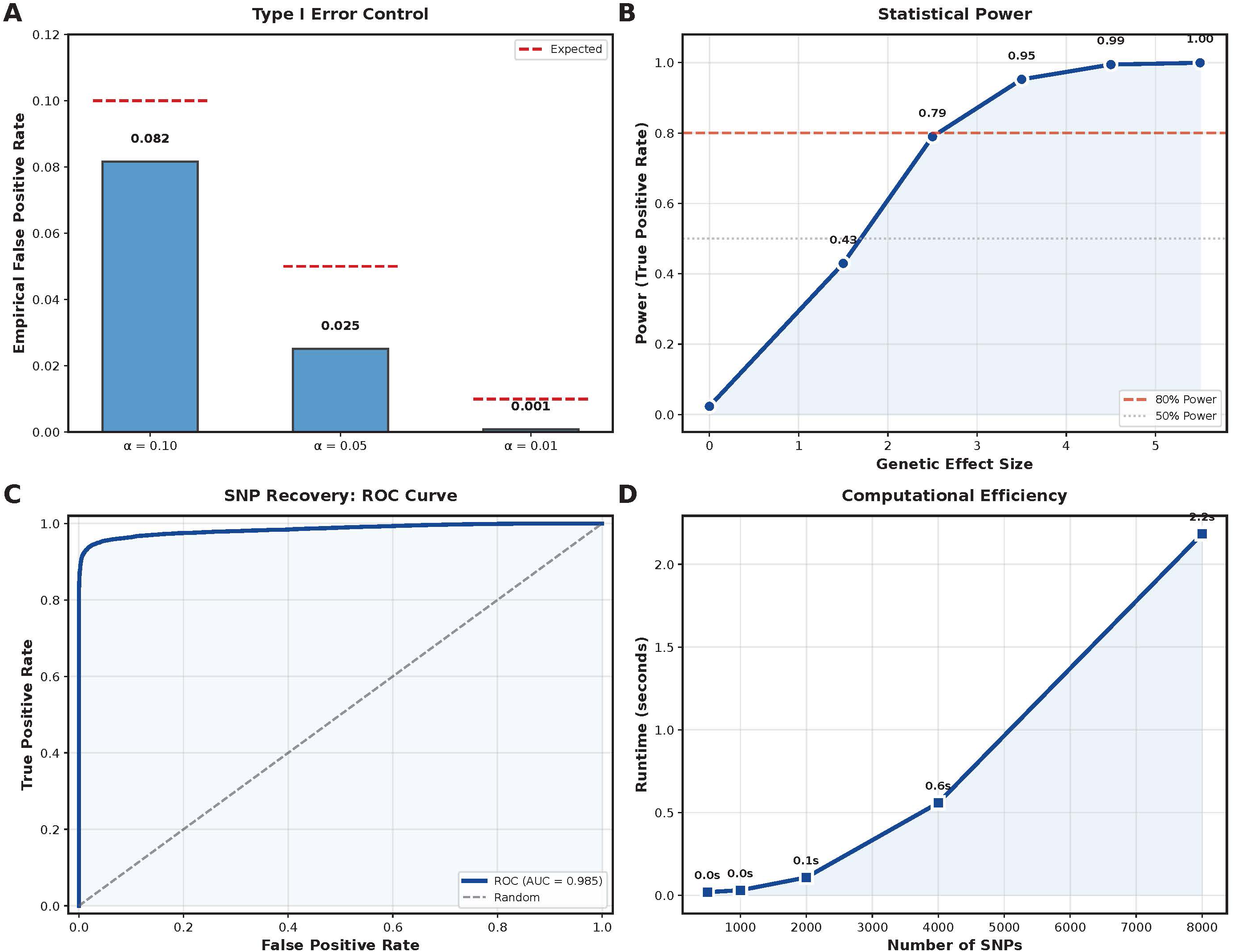
Simulation assessment of PRISMA calibration, recovery, and computational scalability. a, Type I error control. Empirical false positive rates (blue bars) remain below the expected nominal significance levels (𝛼 = 0.10, 0.05, 0.01, red dashed lines), indicating conservative Type I error control against spurious factor discovery. b, Statistical power. The framework showed signal-dependent recovery of injected signal trajectories, approaching the 80% power threshold at signal level 2.5 (78.96%) and surpassing it at signal level 3.5 (95.25%), with high recovery at larger signal magnitudes. c, SNP recovery precision. The ROC curve showed high discriminative performance for separating injected signal variants from background variants, achieving an AUC of 0.985. d, Computational efficiency. The April 18 simulation benchmark shows efficient block-size runtime scaling for the graph-regularized block-wise ALS solver (2,000 SNPs in 0.11 seconds and 8,000 SNPs in 2.18 seconds on an Intel Core i9 processor). Full real-data runtime is reported in Supplementary Table 22.

### Rank selection balances model consistency and biological interpretability

To determine the optimal number of latent axes (rank 𝑅), we evaluated PRISMA across 𝑅 ∈ {1, 2, 3, 4, 5} using CORCONDIA and variance explained metrics (Fig. 3a-b, Supplementary Table 13). Rank 3 satisfied the CORCONDIA criterion (88.1% > 80%) while explaining 85.0% of variance, representing a balanced solution that avoids both underfitting and overfitting. Independent single-cell transcriptomic analyses supported that R=3 corresponds to three axes with distinct tissue-specific cellular enrichment patterns (detailed below), supporting the data-driven rank selection. We therefore selected 𝑅 = 3 as the optimal rank, balancing statistical rigor, parsimony, and biological interpretability.

**Fig. 3.**
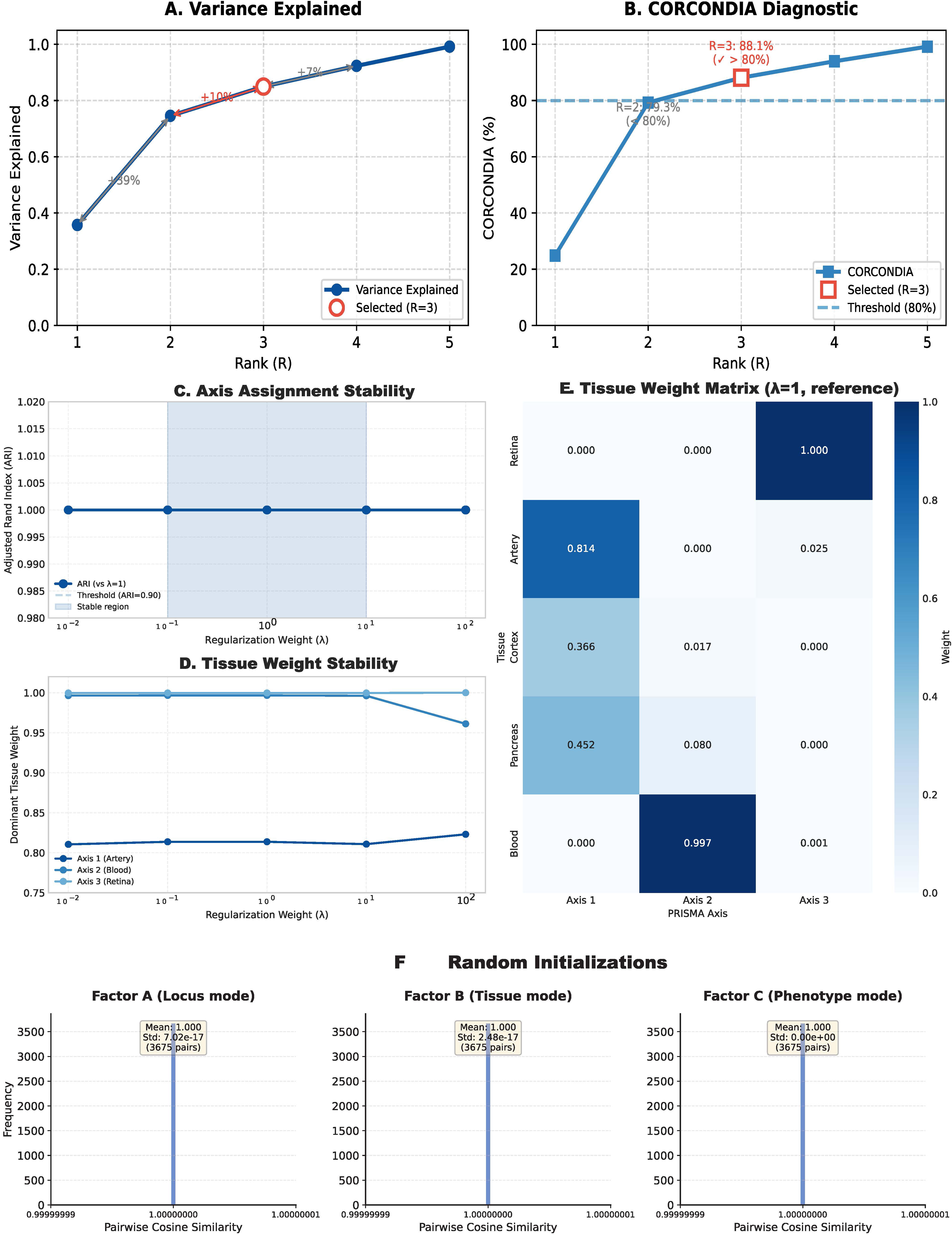
Algorithmic stability and model selection. a-b, Data-driven rank determination. a, Cumulative variance explained across candidate ranks (𝑅 = 1 to 5). b, CORCONDIA. The final rank (𝑅 = 3) was selected as the lowest rank satisfying CORCONDIA > 80% while capturing the major variance-explained elbow and preserving biological interpretability; higher ranks were treated as candidate overfitting regimes requiring additional biological justification. c-e, Sensitivity to topological regularization. c, The ARI of axis-to-tissue assignments was unchanged (ARI = 1.00) across a 10,000-fold perturbation in the graph Laplacian penalty weight (𝜆). d, Dominant tissue weights for the three axes showed limited variation (maximum relative deviation < 3.8%). e, Tissue factor matrix at the 𝜆 = 1 alignment reference used for sensitivity analysis; the production setting (𝜆 = 0.1) yielded the same axis identities. f, Empirical assessment of solution stability. Pairwise cosine similarities of Factor A, B, and C across 50 random initializations showed narrow distributions (mean = 1.000; variance at machine-epsilon scale, ∼ 10^−17^), supporting reproducible convergence under these settings without implying formal identifiability.

Higher-rank solutions explained more variance but were treated as candidate overfitting regimes unless supported by additional biological structure. This rank-selection strategy therefore prioritized a parsimonious decomposition that retained major signal structure while avoiding near-reconstruction solutions driven primarily by numerical fit.

This rank choice was used for the primary DR analysis before downstream biological projection. Subsequent single-cell, baseline, height-control, and vitreous analyses therefore evaluated a pre-specified three-axis solution rather than selecting axes post hoc from the strongest validation signals. This separation helps keep model selection anchored to tensor consistency and variance structure, with downstream analyses used as orthogonal biological support.

### PRISMA deconstructs DR into systemic vascular-metabolic, circulating immune, and retina-specific neurodegenerative trajectories

We next applied PRISMA to deconstruct the multi-trajectory genetic architecture of diabetic retinopathy (DR). Driven by genome-wide summary statistics and targeted multi-tissue eQTLs, PRISMA resolved the DR polygenic signal into three non-orthogonal genetic axes, yielding a pattern consistent with systemic-to-local organization of DR-associated genetic risk. Mapping axis-specific weights back to the genome prioritized 549 top targets, including 403 (73%) below the conventional genome-wide significance threshold (Fig. 4, Supplementary Table 5). These targets should be interpreted as PRISMA-prioritized candidates for tissue-aware follow-up rather than newly established causal loci.

**Fig. 4.**
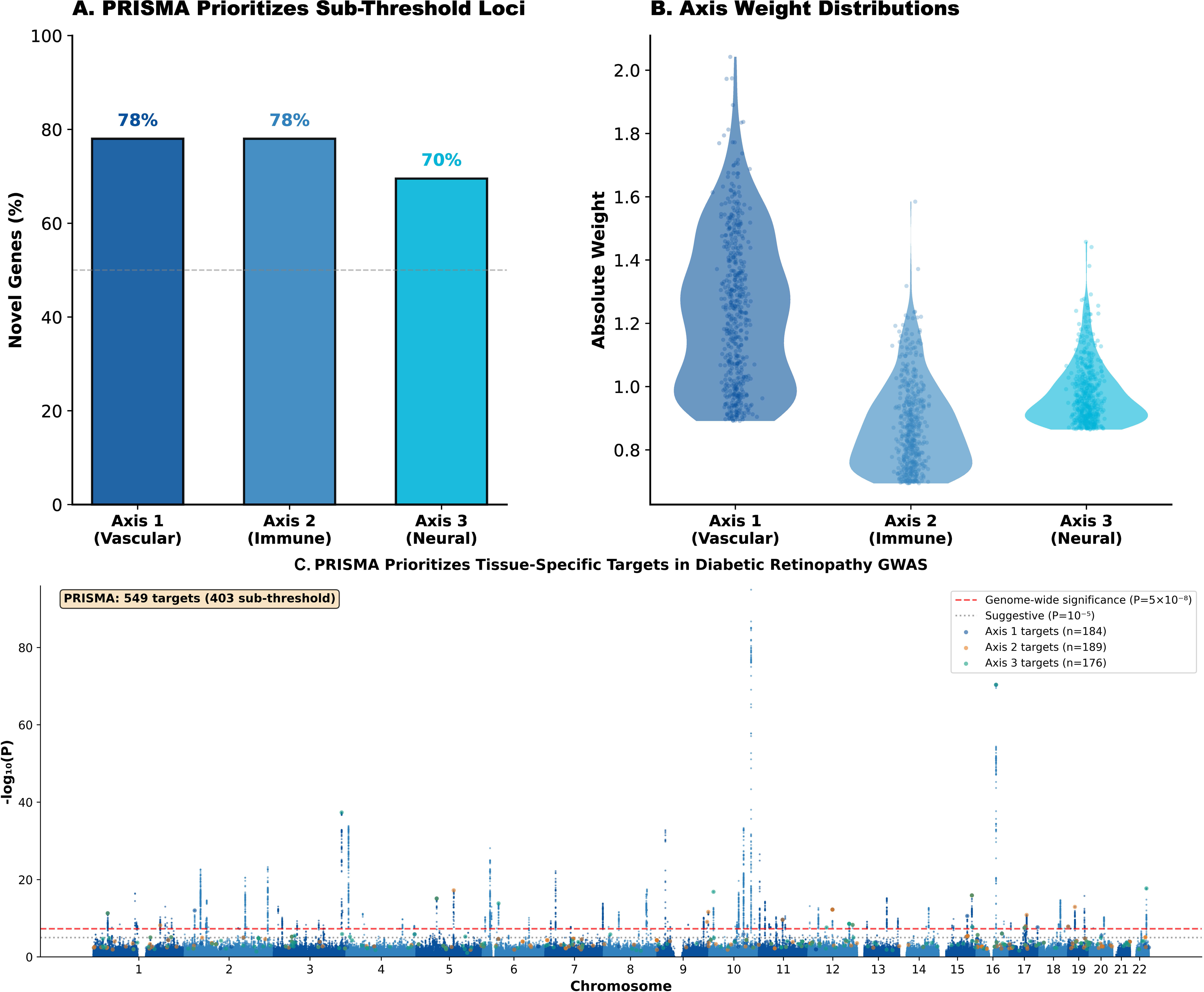
Deconvolution of diabetic retinopathy genetic architecture and prioritization of tissue-specific sub-threshold targets. a, Proportion of PRISMA-prioritized axis-associated genes. Across the three derived axes, approximately 68%-76% of top-weighted targets do not reach conventional genome-wide significance in the baseline GWAS. b, Distribution of absolute factor weights for prioritized targets across the Vascular (Axis 1), Immune (Axis 2), and Neural (Axis 3) trajectories. c, Dual-layer Manhattan plot showing the genome-wide projection of PRISMA-derived targets. Background points represent standard unadjusted GWAS *P*-values for diabetic retinopathy. Colored highlights denote 549 top PRISMA-prioritized targets by genetic risk axis; 403 localize below the conventional genome-wide significance threshold (*P* = 5 × 10^−8^), illustrating the framework’s ability to prioritize sub-threshold polygenic signals and map them to tissue-regulatory pathways.

Axis 1 delineates a “Systemic Vascular-Metabolic Axis”. Tissue-specific factor loadings (Factor B; Supplementary Table 12) showed predominant artery (weight = 0.810) and pancreas (weight = 0.448) anchoring. Projection onto PBMC and proliferative fibrovascular membrane (FVM) single-cell atlases showed spatial divergence (Fig. 5, Supplementary Tables 8 and 15; Supplementary Data 4). Peripheral T cells and platelets showed modest systemic depletion in PBMC, whereas within-FVM cell-type/module-score contrasts showed positive Axis 1 shifts in phagocytic macrophages (*P* = 6.56 × 10^−33^), IFN+ pro-inflammatory macrophages (*P* = 4.61 × 10^−4^), and fibroblasts (*P* = 1.73 × 10^−18^; Supplementary Table 15). In the direction-aware permutation analysis, fibroblasts were the strongest FVM-enriched Axis 1 compartment (*P*_𝑒𝑚𝑝𝑖𝑟𝑖𝑐𝑎𝑙_ = 0.013), while phagocytic and IFN+ macrophages showed positive but weaker trends (*P*_𝑒𝑚𝑝𝑖𝑟𝑖𝑐𝑎𝑙_ = 0.081 and 0.092; Supplementary Table 8).

**Fig. 5.**
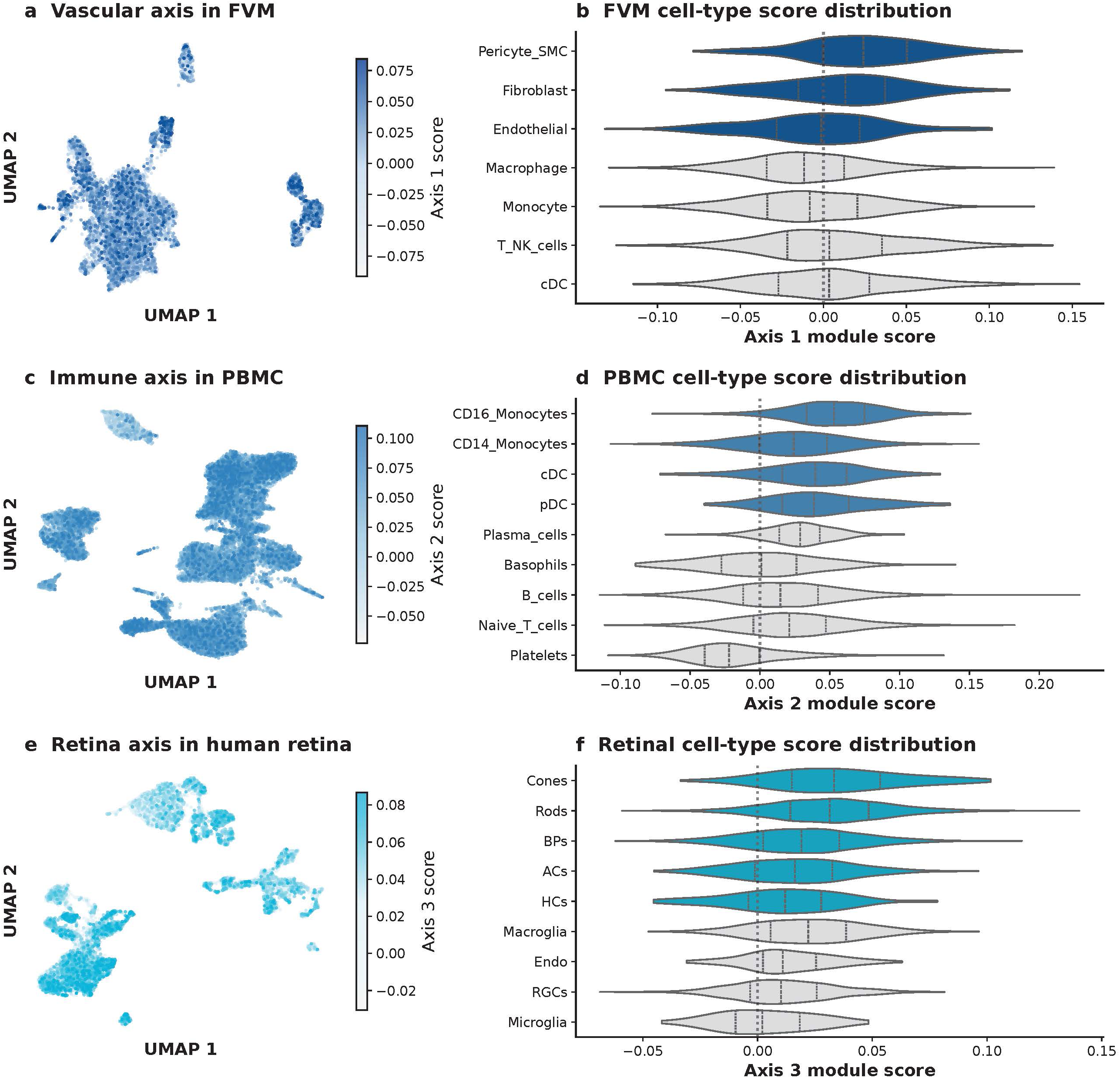
Single-cell spatial mapping of PRISMA latent axes using cell-level module scores. a-b, FVM projection of the vascular-metabolic axis (Axis 1). a, UMAP projection of FVM single cells colored by Axis 1 module scores. b, Cell-type score distributions highlighting Axis 1 signal in pericyte/smooth-muscle and fibroblast compartments, with endothelial, macrophage, and other immune backgrounds shown for comparison. c-d, PBMC projection of the circulating immune-inflammatory axis (Axis 2). c, UMAP projection of PBMC single cells colored by Axis 2 module scores. d, Cell-type score distributions highlighting monocyte and dendritic-cell compartments, consistent with direction-aware permutation results in Supplementary Table 8. e-f, Retinal projection of the retina-specific neurodegenerative axis (Axis 3). e, UMAP projection of the GSE137537 retinal atlas colored by Axis 3 module scores. f, Cell-type score distributions showing strongest Axis 3 scores in photoreceptor and neuronal retinal classes. Cell-level source data are provided in Supplementary Data 4.

Axis 2 represents a “Circulating Immune-Inflammatory Axis” with a whole-blood-dominant tissue profile (weight = 0.996). At single-cell resolution, Axis 2 mapped to circulating CD14+ (*P*_𝑒𝑚𝑝𝑖𝑟𝑖𝑐𝑎𝑙_ = 0.038) and CD16+ (*P*_𝑒𝑚𝑝𝑖𝑟𝑖𝑐𝑎𝑙_ = 0.029) monocytes, as well as conventional dendritic cells (*P*_𝑒𝑚𝑝𝑖𝑟𝑖𝑐𝑎𝑙_ = 0.035), in the PBMC cohort. In FVM, this trajectory showed only marginal fibroblast enrichment (*P*_𝑒𝑚𝑝𝑖𝑟𝑖𝑐𝑎𝑙_ = 0.059) and a weaker positive trend in classical dendritic cells (*P*_𝑒𝑚𝑝𝑖𝑟𝑖𝑐𝑎𝑙_ = 0.105), consistent with a predominantly systemic circulating profile.

Axis 3 captures a “Retina-Specific Neurodegenerative Axis” and was localized to retina (weight = 0.999). Single-cell projection indicated a retinal-anchored genetic trajectory associated with depletion-direction signals in systemic immune compartments. In the PBMC cohort, major immune compartments including monocytes, pDCs, and T cell subsets showed depletion-direction shifts. In FVM, Axis 3 showed no positive enrichment by direction-aware permutation testing (all *P*_𝑒𝑚𝑝𝑖𝑟𝑖𝑐𝑎𝑙_ ≥ 0.248), supporting spatial decoupling from the local fibrovascular program. Together, these findings are consistent with genetic heterogeneity that is organized across tissue contexts; however, these correlations do not establish temporal or causal ordering.

The three-axis structure therefore separates vascular-metabolic, circulating immune, and retinal-neural regulatory profiles without requiring those profiles to be mutually exclusive. Because PRISMA factors are non-orthogonal, shared loci and partially overlapping downstream pathways can coexist with tissue-biased axis definitions. This structure is consistent with the broader expectation that complex disease risk may involve both divergence in upstream regulatory routes and convergence in downstream pathology.

Table 1 summarizes the dominant tissue loadings, orthogonal single-cell support, exploratory vitreous patterns, and inference boundaries for the three PRISMA-derived DR axes.

**Table 1.** Summary of PRISMA-derived diabetic retinopathy axes. Dominant tissue loadings are from Factor B (Supplementary Table 12). Single-cell support summarizes direction-aware empirical permutation and module-score analyses (Supplementary Tables 8 and 15; Supplementary Data 4). Proteomic module scores were elevated in DR for all three axes; metabolite-pattern rows summarize sensitivity-filtered metabolite-axis associations (Supplementary Tables 9, 11, and 17). Vitreous metabolite specificity was generally low (median *S*_𝑚_ = 0.016), so these molecular patterns should be interpreted as candidate downstream correlates rather than causal mechanisms or stable biomarkers. An expanded editable version of this axis summary is provided in Supplementary Table 23.

| Axis | Trajectory label | Dominant tissue loading | Key orthogonal support and boundary |
| --- | --- | --- | --- |
| Axis 1 | Systemic<br>vascular-metabolic | Artery 0.810; pancreas<br>0.448 | FVM fibroblast<br>enrichment<br>($P_{\text{empirical}} = 0.013$);<br>29/35 filtered<br>metabolites.<br>Exploratory and<br>non-causal. |
| Axis 2 | Circulating<br>immune-inflammatory | Whole blood 0.996 | PBMC<br>monocyte/dendritic-cell<br>enrichment<br>($P_{\text{empirical}} = 0.029$-<br>0.038); 4/35 filtered<br>metabolites.<br>Exploratory and<br>non-causal. |
| Axis 3 | Retina-specific neurodegenerative | Retina 0.999 | Retinal neuronal enrichment ($P_{\text{empirical}} < 0.001$), with no positive FVM enrichment; 2/35 filtered metabolites. Exploratory and non-causal. |

### Spatial decoupling and double dissociation of genetic risk axes

To evaluate whether PRISMA-derived axes show spatially distinct enrichment beyond generic disease-gene signals, we performed cross-axis permutation tests. For each axis, 200 genes were randomly sampled from the background pool of DR-associated genes input to PRISMA, repeated 1,000 times, and cell-type module scores were recomputed across PBMC and FVM datasets. This null distribution tests whether PRISMA gene sets show stronger tissue-specific enrichment than random disease-gene sets of equal size.

The results showed a double-dissociation pattern (Fig. 6, Supplementary Tables 8 and 15). Axis 1 exhibited significant enrichment in FVM fibrovascular cells (*P*_𝑒𝑚𝑝𝑖𝑟𝑖𝑐𝑎𝑙_ = 0.013) but not in PBMC cell types (all *P*_𝑒𝑚𝑝𝑖𝑟𝑖𝑐𝑎𝑙_ > 0.17). Axis 2 showed enrichment in PBMC monocytes and dendritic cells (*P*_𝑒𝑚𝑝𝑖𝑟𝑖𝑐𝑎𝑙_ < 0.04) but only marginal significance in FVM fibroblasts (*P*_𝑒𝑚𝑝𝑖𝑟𝑖𝑐𝑎𝑙_ = 0.059). In contrast, Axis 3 showed strong enrichment in retinal neurons (photoreceptors, bipolar cells: all *P*_𝑒𝑚𝑝𝑖𝑟𝑖𝑐𝑎𝑙_ < 0.001) but no positive enrichment in FVM (all *P*_𝑒𝑚𝑝𝑖𝑟𝑖𝑐𝑎𝑙_ ≥ 0.248) or PBMC; significant PBMC signals, when present, were depletion-direction effects with negative Δ Mean (Supplementary Table 8). These results support spatially biased enrichment patterns consistent with expected tissue contexts. A permutation sensitivity analysis across sliding cutoffs (Top 50 to all 243 strictly Axis 1 dominant targets) showed that Axis 1 FVM enrichment remained significant across thresholds (*P* ≤ 0.01, Supplementary Fig. 3, Supplementary Table 16), indicating that axis weights are not explained by arbitrary thresholding alone.

**Fig. 6.**
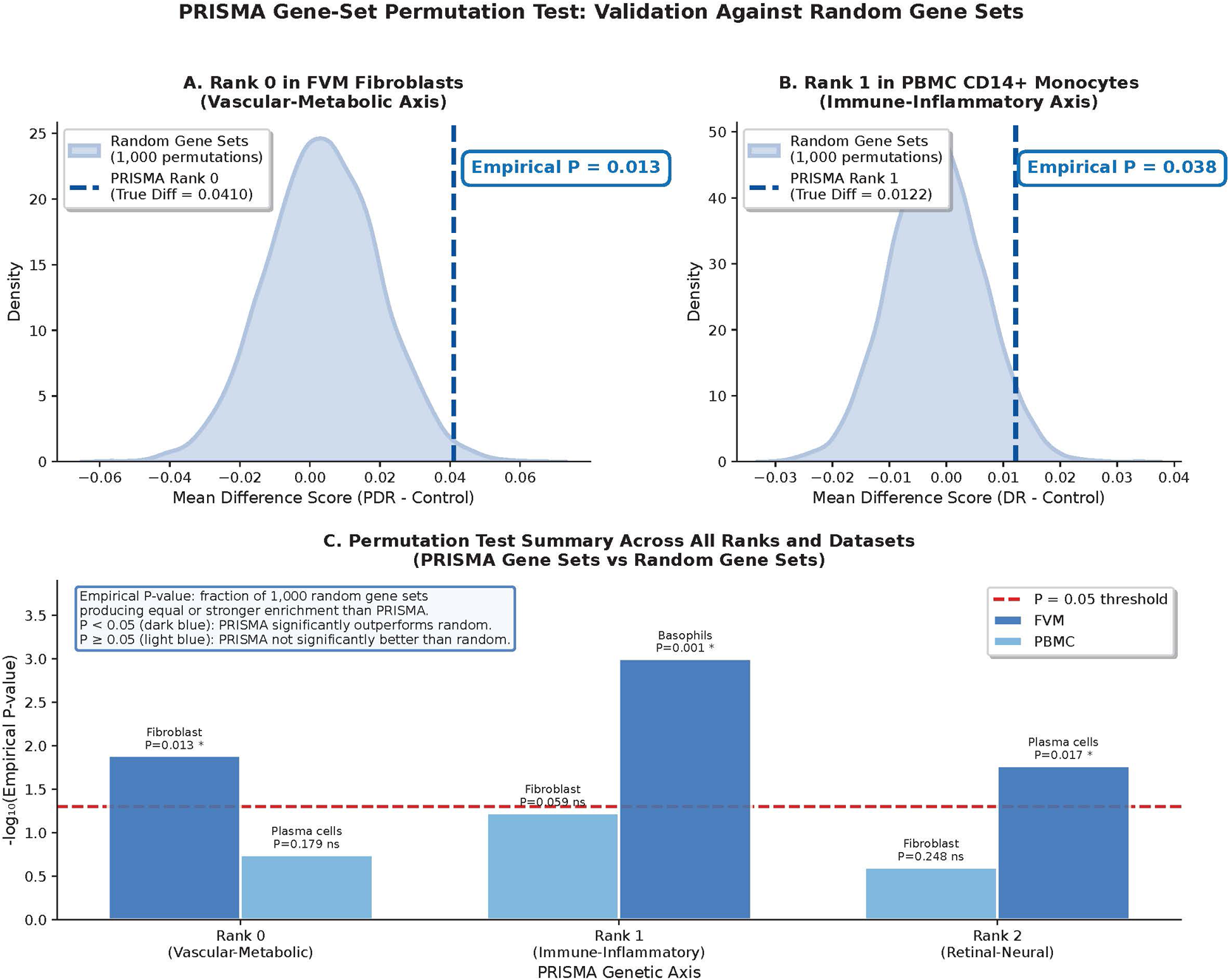
Gene-set permutation test for single-cell trajectory enrichment. Empirical null distributions were generated from 1,000 random permutations using background disease-associated gene sets of identical size to evaluate cell-type module scores. a-b, Representative null distributions (shaded areas) of module-score mean differences for Axis 1 in FVM fibroblasts (a) and Axis 2 in PBMC CD14+ monocytes (b). Dashed vertical lines indicate observed effect sizes of PRISMA-derived gene sets and empirical P-values. c, Summary of permutation results across the three axes and tested single-cell microenvironments. Bar heights represent − log_10_(empirical *P*-value). The red dashed line denotes nominal 𝛼 = 0.05. Dark blue bars highlight significant direction-aware deviations; positive-enrichment and depletion-direction effects are distinguished in Supplementary Table 8.

These permutation analyses are important because they compare PRISMA gene sets against random draws from the same disease-associated candidate background rather than against an unrestricted transcriptome. The tests therefore address whether the axes encode cell-type specificity beyond generic DR gene membership, while still avoiding causal interpretation of the observed module-score differences.

### PRISMA characterizes biologically meaningful tissue architectures missed by conventional dimensionality reduction

To evaluate whether these tissue-specific trajectories arise from PRISMA’s LD-aware tensor framework rather than generic dimensionality reduction, we benchmarked PRISMA against PCA, NMF, and K-means applied to the identical PRISMA integration-score matrix (8,201 SNPs × 5 tissues). All methods extracted 3 components matching PRISMA’s rank. Tissue specificity was quantified using the Gini coefficient (0 = uniform distribution, 1 = single-tissue dominance). PRISMA produced higher tissue specificity (Gini = 0.688 ± 0.129) than NMF (0.272 ± 0.150), PCA (0.233 ± 0.129), or K-means (0.147 ± 0.076) (Supplementary Fig. 1, Supplementary Table 7).

This comparison supports that PRISMA produced sharper tissue-specific profiles than the tested baselines. The difference is consistent with the graph Laplacian regularization, which preserves local LD structure and favors LD-coherent regulatory patterns. By contrast, PCA, NMF, and K-means lack this genomic-topology constraint and showed weaker alignment with interpretable tissue-regulatory architectures. Having evaluated whether PRISMA identifies tissue-specific patterns not captured by conventional methods, we next examined whether these axes were associated with molecular signatures in the disease microenvironment.

The baseline analysis also clarifies the role of non-negativity alone. NMF imposes non-negative factors but does not encode local LD topology, and it produced substantially more diffuse tissue profiles than PRISMA. This suggests that the combination of non-negativity with graph-based LD regularization, rather than non-negativity by itself, is central to the observed tissue resolution.

### Established post-GWAS comparators support gene-level concordance of PRISMA-prioritized targets

Because the 549 PRISMA target rows corresponded to 405 unique gene symbols, established gene-level comparator analyses were performed using this 405-gene projection (Supplementary Table 24). MAGMA showed significant GWAS gene-set burden for the PRISMA target set (BH FDR = 2.89 × 10^−5^) and for all three axis-specific gene sets, including the exploratory subthreshold set (Supplementary Table 25). Complementary SMR analysis showed that strict SMR-supported genes were enriched among PRISMA-prioritized genes within the PRISMA background universe (31 genes; OR = 6.32, *P* = 1.95 × 10^−13^; Supplementary Table 26). Finally, as a targeted shared-variant sensitivity analysis, coloc.abf was applied only to QC-passed strict SMR-PRISMA gene-tissue pairs; 12 of 23 eligible pairs reached PP.H4 ≥ 0.80, corresponding to 9 of 16 tested genes, with another 5 pairs showing suggestive support (Supplementary Table 27). Together, these comparator analyses support gene-level concordance between PRISMA prioritization and established post-GWAS evidence, without implying causal validation of all PRISMA targets.

### Height GWAS negative control supports trait-dependent regulatory reprioritization

To test whether PRISMA mainly captures generic tissue expression architecture, we applied the identical tensor framework to adult height GWAS summary statistics from GWAS Catalog accession ebi-a-GCST90018959 (European ancestry, N=360,388) [12,13], using the same five-tissue eQTL panel. Height is highly polygenic and uses overlapping regulatory architecture but has a distinct biological relationship to DR.

PRISMA decomposed the height tensor into three tissue-specific axes. In this matched negative-control rerun, height and DR tensors showed comparable model-fit and tissue-segregation summaries (Height: CORCONDIA = 64.6%, mean Gini = 0.687; matched DR rerun: CORCONDIA = 64.6%, mean Gini = 0.688; Supplementary Fig. 2, Supplementary Tables 6 and 19), supporting transferability of the optimization framework without conflating this rerun with primary DR rank selection. Matching decomposed ranks by primary tissue loading showed substantial top-200 gene overlap (39-45%) relative to the shared eQTL-mappable gene universe (N = 8,177; hypergeometric test, all *P* < 10^−77^; Supplementary Table 19). This overlap reflects a shared tissue-regulatory backbone rather than disease-specific equivalence. Trait-dependent reprioritization appeared in relative weights: DR-weighted genes such as TAGLN and ALDH16A1 were present in the height gene pool but ranked lower, while height-prioritized examples such as GBP3 and CUTALP showed the inverse pattern (Supplementary Table 19). Collectively, these results support trait-dependent regulatory prioritization within a shared eQTL-mappable tissue framework.

Thus, the negative control did not show that DR and height use disjoint regulatory genes; instead, it showed that a shared eQTL-mappable background can be reweighted by distinct GWAS signals. This is the expected behavior for a polygenic regulatory model and supports the interpretation that PRISMA captures relative prioritization within a common regulatory search space.

### Exploratory vitreous multi-omics analysis suggests candidate molecular correlates of PRISMA axes

To explore whether PRISMA genetic trajectories associate with molecular signatures, we performed an exploratory paired multi-omics analysis of vitreous humor from 13 patients (7 with severe proliferative DR and 6 non-diabetic controls). Proteomics and metabolomics were treated as hypothesis-generating down-stream analyses. PRISMA module scores were elevated in DR compared to controls (Supplementary Table 17; Mann-Whitney U test: Axis 1, *P* = 1.17 × 10^−3^; Axis 2, *P* = 1.17 × 10^−3^; Axis 3, *P* = 1.17 × 10^−3^).

Metabolite-axis correlation analysis (Fig. 7, Supplementary Tables 9 and 11) suggested a pattern compatible with convergence of genetically separable PRISMA trajectories onto a shared vitreous metabolic milieu rather than fully independent biochemical modules. In exploratory FDR analysis, metabolite-axis correlations passing q < 0.05 were observed for all three trajectories (Axis 1, n = 125; Axis 2, n = 83; Axis 3, n = 117; Supplementary Table 11). After FDR, effect-size, and leave-one-out cross-validation filtering, 35 metabolite features passed exploratory sensitivity filters (Supplementary Table 9), with Axis 1 predominance (29 Axis 1, 4 Axis 2, and 2 Axis 3 features) and generally low orthogonal specificity (median *S*_𝑚_ = 0.016). The retained Axis 1-associated features were mainly lipid and membrane-remodeling features, while Axis 2 and Axis 3 had smaller filtered subsets.

**Fig. 7.**
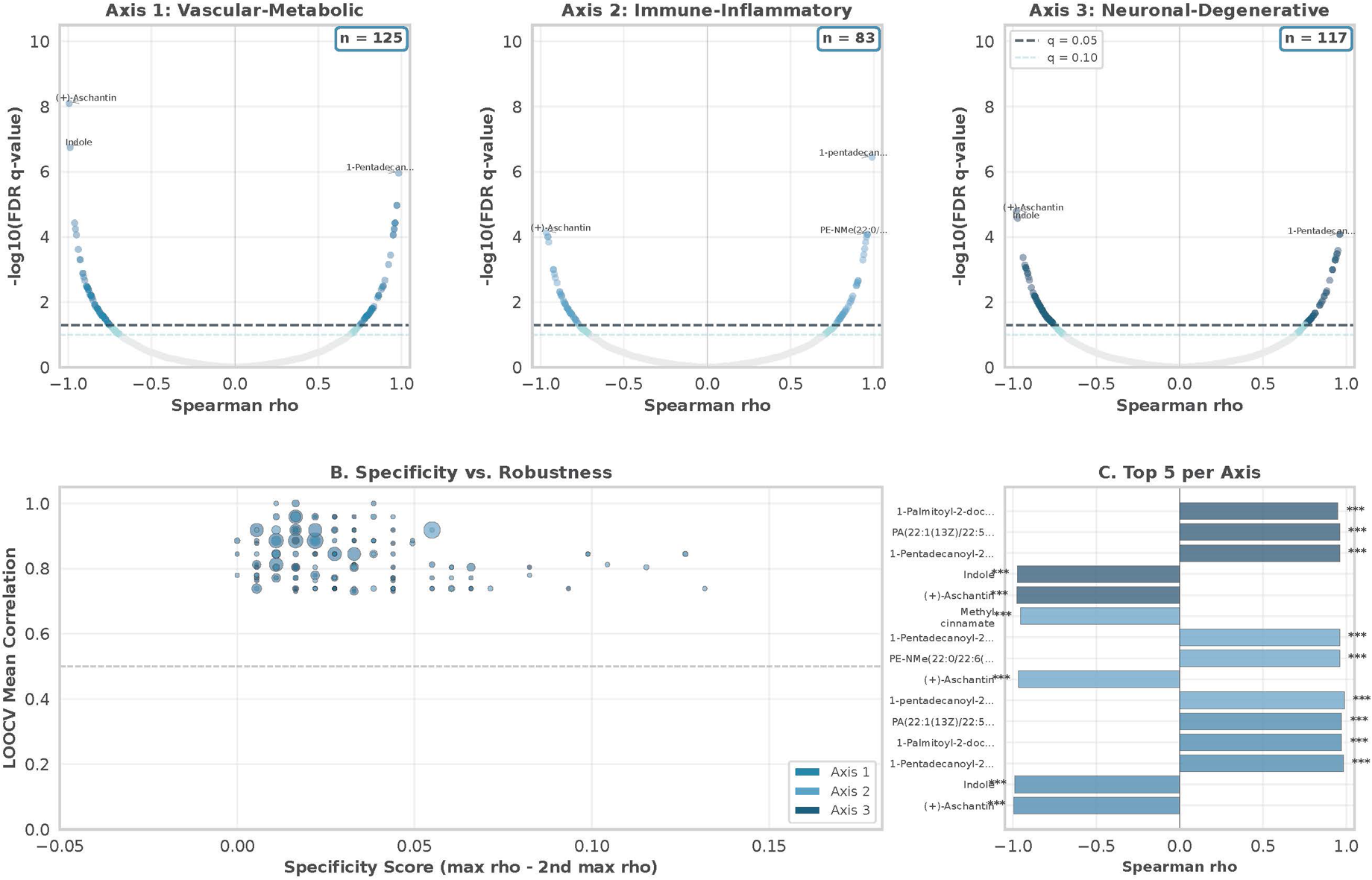
Exploratory vitreous metabolomic analysis and candidate molecular convergence patterns. a, Volcano-style plots showing correlations between individual-level PRISMA module scores and vitreous metabolite abundances across the Vascular-Metabolic (Axis 1), Immune-Inflammatory (Axis 2), and Neuronal-Degenerative (Axis 3) trajectories. The U-shaped distribution reflects the deterministic relationship between Spearman’s rank correlation (𝜌) and FDR q-value at fixed sample size (n=7 severe DR patients). Dashed lines indicate q=0.05 and q=0.10. b, Distribution of metabolite trajectory specificity versus cross-validation robustness. Clustering of significant metabolites at low specificity scores is compatible with, but does not establish, downstream convergence. c, Top FDR-ranked metabolite-axis associations per axis, highlighting weak axis-linked signals embedded within a shared vitreous metabolic milieu. Source data are provided in Supplementary Data 2. Given the limited sample size, these molecular associations are exploratory and hypothesis-generating.

The low median specificity score is notable because it indicates that many significant features were not cleanly assigned to only one genetic axis. Rather than treating this as evidence against the axis model, we interpret it cautiously as a molecular pattern compatible with partial convergence in bulk vitreous fluid. In this anatomical compartment, downstream extracellular signals may integrate vascular leakage, inflammatory activity, and neuroretinal stress, reducing the apparent specificity of individual metabolite features.

These results are qualitatively compatible with an upstream-divergence/downstream-convergence model: spatially distinct genetic axes retain weak axis-linked molecular signatures but converge onto a shared vitreous metabolic environment dominated by vascular-metabolic and lipid-remodeling processes. However, the metabolomics correlations were restricted to n=7 DR patients and do not establish causal directionality, temporal ordering, stable biomarkers, or predictive value. These molecular associations should therefore be interpreted as candidate correlates of PRISMA axes rather than stable biomarkers. Larger longitudinal cohorts with matched genotype and molecular profiling will be needed to test temporal ordering and mechanistic convergence.

## Discussion

PRISMA was developed to address a specific limitation of conventional complex-disease GWAS interpretation: heterogeneous, tissue-dependent regulatory effects are usually compressed into global locus-level statistics. By integrating GWAS summary statistics with multi-tissue cis-eQTL data and constraining factorization by local LD topology, PRISMA provides a quantitative framework for resolving tissue-biased genetic trajectories from aggregate association signals. Its contribution is therefore not simply to nominate individual genes, but to quantify genetic heterogeneity that is already present in GWAS and eQTL data yet obscured by tissue-collapsed analysis. In the present diabetic retinopathy application, the framework was instantiated in the single-phenotype setting, where the phenotype mode is a singleton dimension and Factor C acts as a rank-specific scaling vector. Within this single-phenotype setting, the empirical contribution of PRISMA is an LD-aware, summary-statistics decomposition of locus-by-tissue genetic architecture, establishing the single-trait foundation for future multi-phenotype tensor applications.

Applied to DR, PRISMA resolved the aggregated polygenic signal into three tissue-biased axes with interpretable biological structure: a vascular-metabolic axis anchored in artery and pancreas, a circulating immune-inflammatory axis dominated by whole-blood regulation, and a retina-specific neurodegenerative axis. These axes align with major biological themes in DR while refining them at the tissue-regulatory level. The vascular-metabolic axis is consistent with classical metabolic and microvascular mechanisms, including pathways linked to hyperglycemic injury and vascular barrier dysfunction [14–17]. The immune-inflammatory axis is consistent with systemic inflammatory, immune-remodeling, and monocyte-linked models of diabetic microvascular complications [18–23]. The retina-specific axis supports the view that DR includes a neuroretinal component rather than being exclusively microvascular [24]. These interpretations should be read as genetically anchored regulatory hypotheses rather than direct mechanistic proof, because the model operates on summary-level GWAS and eQTL evidence.

A central feature of PRISMA is that LD topology is incorporated during decomposition rather than appended as a post hoc annotation. Variants within local haplotypic neighborhoods are not independent observations; they share physical co-inheritance structure that can make unconstrained matrix factorization unstable or biologically diffuse. The graph Laplacian penalty encourages LD-connected variants to carry coordinated latent weights, thereby anchoring the factorization to local genomic structure while retaining tissue-level contrast. This design helps explain why PRISMA produced sharper tissue-regulatory profiles than PCA, NMF, or K-means applied to the same SNP-by-tissue matrix.

Orthogonal single-cell analyses supported the tissue interpretation of the PRISMA axes. Axis 1 was enriched in local fibrovascular compartments, Axis 2 was enriched in circulating monocyte and dendritic-cell compartments, and Axis 3 showed retinal neuronal enrichment with no positive fibrovascular enrichment. This spatial double-dissociation provides biological support that the axes are not simply generic disease-gene sets or tissue expression artifacts. The height GWAS negative-control analysis further supported trait-dependent reprioritization: using the same eQTL panel, PRISMA recovered a shared eQTL-mappable tissue backbone but shifted gene weights according to the GWAS trait. Together, these analyses suggest that PRISMA captures regulatory prioritization shaped by disease-associated genetic signal rather than merely clustering ubiquitous eQTL architecture.

The exploratory vitreous proteomic and metabolomic analyses provide a separate downstream perspective on the inferred genetic axes. The observation that genetically separable axes mapped to overlapping vitreous molecular correlates is compatible with an upstream-divergence and downstream-convergence model: distinct tissue-regulatory routes may contribute to a shared pathological molecular environment in advanced DR. This interpretation is useful because it explains why upstream genetic heterogeneity may coexist with clinically convergent disease manifestations. However, it remains hypothesis-generating. The metabolomics correlation analysis was restricted to seven DR samples, and the observed correlations cannot establish temporal order, causal direction, or stable biomarkers. The convergence model should therefore be viewed as an organizing hypothesis that motivates larger longitudinal multi-omics studies rather than as a definitive mechanistic conclusion.

Methodologically, PRISMA differs from both aggregate heritability frameworks and generic dimensionality-reduction approaches. S-LDSC and related methods estimate global enrichment across annotations but do not assign disease-associated loci to tissue-biased trajectories. Fine-mapping methods prioritize causal variants but generally do not decompose a single disease into latent locus-by-tissue regulatory axes. PCA, NMF, and K-means can summarize variation in a SNP-by-tissue matrix, but they do not incorporate LD topology during decomposition. PRISMA instead regularizes locus-level factor loadings by an empirical LD graph and interprets the resulting axes as tissue-resolved genetic trajectories. In this sense, PRISMA complements rather than replaces existing GWAS interpretation tools: it provides a decomposition layer between locus discovery and downstream biological validation.

The post-GWAS comparator analyses empirically support this complementary positioning. After projecting the 549 PRISMA-prioritized target rows to 405 unique gene symbols, the resulting gene set showed significant MAGMA burden, enrichment for strict SMR/HEIDI-supported genes, and targeted colocalization support for a subset of eligible SMR-PRISMA gene-tissue pairs. These findings indicate that PRISMA-prioritized targets are concordant with established gene-level GWAS-eQTL evidence, while PRISMA retains a distinct output: the organization of those targets into LD-aware, tissue-biased genetic trajectories. Importantly, these comparator analyses should not be interpreted as causal validation of all PRISMA targets. Rather, they show that PRISMA operates within the evidentiary landscape captured by established post-GWAS tools while adding a trajectory-level decomposition layer.

The prioritization of 549 axis-associated targets, including many below conventional genome-wide significance thresholds, should be interpreted in this framework. These targets are not claimed as newly established causal loci. Rather, PRISMA uses regulatory and topological structure to nominate sub-threshold polygenic signals for tissue-aware follow-up. This distinction is important for complex diseases, where meaningful polygenic architecture may be distributed across many loci that individually fall below strict discovery thresholds but collectively define coherent biological trajectories. Clinical observations of heterogeneous DR phenotypes and variable therapeutic contexts provide additional motivation for treating these axes as hypotheses for future stratification rather than validated endotypes [25–28].

The use of summary-level data is an important practical feature. PRISMA does not require individual-level genotype, transcriptome, or phenotype data for its core decomposition, making it deployable across diseases for which GWAS and eQTL summary statistics are available. This accessibility is particularly relevant for diseases where disease-relevant tissues are difficult to sample, such as retina, and it supports broader reuse once code and documentation are publicly released with the preprint. At the same time, summary-level accessibility imposes constraints. PRISMA cannot by itself determine whether a prioritized regulatory trajectory is causal, compensatory, or secondary to disease progression. Individual-level genotypes, longitudinal clinical phenotypes, perturbation experiments, and ideally disease-relevant molecular QTLs will be required to test the biological and clinical implications of the inferred axes.

Although PRISMA is formulated as a general locus-by-tissue-by-phenotype tensor framework, the present study deliberately instantiated the model in the single-phenotype setting. In this setting, the phenotype mode is fixed at *P* = 1, and Factor C acts as a rank-specific scaling vector rather than a multi-trait loading matrix. This design was chosen to enable deep biological evaluation of tissue-resolved genetic trajectories within one clinically heterogeneous complex disease, diabetic retinopathy, using orthogonal single-cell transcriptomic atlases and an exploratory clinical vitreous multi-omics cohort. The retained 𝑁 × 𝑇 × *P* formulation defines the mathematical basis for multi-phenotype applications, but the present study does not empirically evaluate the *P* > 1 setting. Analyses in that setting would require additional treatment of cross-trait dependence, phenotype nesting, and potential sample overlap among GWAS summary statistics.

Several limitations should be emphasized. First, as noted above, the current empirical analysis is restricted to one phenotype, DR, despite the more general tensor formulation. Second, tissue selection was hypothesis-driven and limited to five DR-relevant tissues; tissues or cell states without eQTL coverage could not contribute to the decomposition. Third, the single-cell atlases were used for orthogonal enrichment analyses rather than causal validation, and the retinal atlas was not a DR-specific perturbation dataset. Fourth, the vitreous multi-omics cohort was small, especially for metabolomics correlations among DR patients, and should be treated as exploratory. Fifth, although FinnGen and MVP provided substantial GWAS sample size, cohort selection effects can influence representation of complication-rich phenotypes [29], and independent replication of the inferred axes in additional DR cohorts and across ancestries will be needed to evaluate robustness and generalizability.

In summary, PRISMA reframes complex-disease GWAS from aggregate locus discovery toward quantitative, LD-aware mapping of tissue-resolved genetic trajectories. In DR, this approach organized the polygenic signal into vascular-metabolic, immune-inflammatory, and retina-specific neurodegenerative trajectories and nominated axis-associated targets for tissue-aware follow-up. More broadly, PRISMA positions summary-statistics genetics as a reusable decomposition layer between GWAS discovery and experimental functional validation: it does not replace causal inference or mechanistic experiments, but it can prioritize where such experiments should look. The present study provides proof-of-principle evidence for trajectory-level genetic dissection using summary statistics, while leaving causal validation, prospective stratification testing, and multi-phenotype extension as necessary next steps.

## Methods

### Ethics statement

The acquisition and use of human vitreous humor samples was approved by the Institutional Review Board of the Eye Hospital of China Academy of Chinese Medical Sciences, Beijing. Approval number: YKEC-KT-2024-054-P003. Written informed consent was obtained from all participants. Publicly available summary-level data (FinnGen, MVP, GTEx, GEO, and GWAS Catalog resources) did not require additional ethical approval.

### GWAS, eQTL, and LD reference data

Diabetic retinopathy (DR) GWAS summary statistics were obtained from two European-ancestry resources: the FinnGen research project (R12) [30] and the Million Veteran Program (MVP) [31,32] (Supplementary Table 1). A fixed-effect inverse-variance weighted meta-analysis was performed using METAL [33], yielding a combined sample size of N = 505,435. Standard GWAS quality-control filters removed low-frequency, low-imputation-quality, strand-ambiguous, and invalid standard-error variants; genomic coordinates were referenced to GRCh37 and aligned to the 1000 Genomes Project Phase 3 European reference panel [34]. Full allele harmonization and filtering details are provided in Supplementary Method 1.

Cis-eQTL summary statistics were assembled for five DR-relevant tissues: retina, tibial artery, brain cortex, pancreas, and whole blood (Supplementary Table 2). Retina-specific eQTLs were obtained from a human retina atlas [35], and tibial artery, brain cortex, pancreas, and whole-blood eQTLs were obtained from GTEx Release v8 [36]. For each gene, the strongest cis-eQTL by cross-tissue composite absolute Z-score was retained as the sentinel instrument. Pre-specified filters removed broadly expressed translation-related genes, known inversion-region genes, and complex LD blocks likely to introduce long-range structural artifacts [37–39]. These harmonization, sentinel-selection, and artifact-filtering steps are detailed in Supplementary Method 1.

This preprocessing produced a harmonized set of disease-associated regulatory instruments suitable for locus-by-tissue decomposition rather than single-variant fine-mapping. The filtering strategy was designed to construct a conservative, interpretable tensor input in which each retained gene is represented by one sentinel cis-eQTL and each variant can be assigned consistently across tissues and LD blocks. This reduces redundancy from highly correlated local instruments while preserving tissue-resolved regulatory contrast.

### GWAS-eQTL integration score

To map locus-level disease associations to tissue-specific regulatory contexts, PRISMA integrates GWAS and eQTL summary statistics using a PRISMA-specific Z-score feature weight. For each aligned variant-gene-tissue entry, the GWAS-eQTL integration score is:

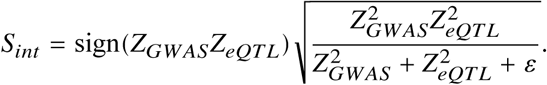

where 𝑍_𝐺𝑊𝐴*S*_ is the disease association Z-score, 𝑍_𝑒𝑄𝑇𝐿_ is the tissue-specific eQTL Z-score, and 𝜀 = 10^−8^ prevents numerical instability. The score assigns large magnitude only when both disease association and local regulatory evidence support the variant-gene-tissue entry. In PRISMA, *S*_*int*_ is used as a feature weight for tensor factorization rather than as formal causal SMR evidence; allele sign conventions, numerical edge cases, and the distinction from HEIDI-based causal SMR are described in Supplementary Method 2.

The sign of *S*_*int*_ records concordance or discordance between the GWAS and eQTL Z-scores, while its magnitude emphasizes entries jointly supported by both sources. Thus, a strong GWAS association with weak eQTL evidence, or a strong eQTL with little disease association, is down-weighted relative to entries supported by both layers. This construction provides a summary-statistics bridge between disease association and tissue-specific regulation without requiring individual-level genotype, expression, or phenotype data.

### Tensor construction and LD graph regularization

Following targeted purification, PRISMA organizes *S*_*int*_ values into genomic-block-specific variant-by-tissue matrices:

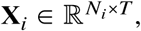

where 𝑁_𝑖_ is the number of variants in LD block 𝑖 and 𝑇 is the number of tissues. The general PRISMA specification is a three-way tensor:

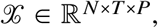

where *P* denotes the phenotype dimension. The present DR analysis evaluates the single-phenotype case (*P* = 1), so the phenotype mode is a singleton dimension and Factor C acts as a rank-specific scaling vector. This makes the present implementation algebraically equivalent to a graph-regularized coupled matrix factorization across genomic blocks while retaining the general tensor notation for future multi-phenotype extensions. Tensor construction details, including sample-overlap considerations for future *P* > 1 analyses, are provided in Supplementary Method 3.

To reduce the influence of isolated extreme Z-scores, PRISMA applies a signed square-root transformation to each block:

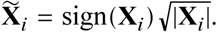

with bounded transformed values. Local LD topology is encoded using the 1000 Genomes Phase 3 European reference panel [34]. For each block, the Pearson correlation matrix **R** among variants is converted to an adjacency matrix:

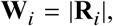

with zero diagonal, and the graph Laplacian is:

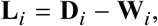

where **D**_𝑖_ is the diagonal degree matrix. The rationale for using |𝑅| rather than 𝑅^2^, and the comparison with rank-based inverse normal transformation, are given in Supplementary Method 3.

The graph Laplacian penalty uses this LD graph to encourage variants in the same local haplotypic neighborhood to have coordinated factor loadings. This is the main distinction between PRISMA and generic decomposition of a flattened SNP-by-tissue matrix: local genomic topology is preserved during factorization rather than treated as a post hoc annotation. The resulting factors can therefore be interpreted as tissue-resolved regulatory trajectories constrained by both GWAS-eQTL evidence and LD structure.

### Graph-regularized block-wise factorization

PRISMA decomposes each genomic block using low-rank factors **A**_𝑖_ ∈ ℝ^𝑁𝑖×𝐾^, **B** ∈ ℝ^𝑇×𝐾^, and **C** ∈ ℝ^1×𝐾^ in the current single-phenotype application. Here, 𝐾 denotes the decomposition rank and is equivalent to 𝑅 used in the rank-selection analysis. **A**_𝑖_ contains locus-level genetic drivers, **B** contains global tissue specificity weights, **C** contains rank-specific phenotype scaling, and 𝐾 is the selected rank. The graph-regularized objective is:

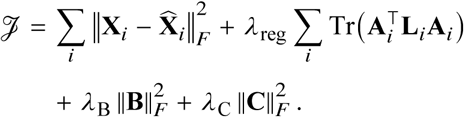

For *P* = 1, **C** reduces to a 1 × 𝐾 scaling vector and the reconstruction simplifies to:

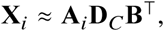

where **D**_𝐶_ is the diagonal matrix formed from **C**. The model is optimized by block-wise alternating least squares [43,44]: local **A**_𝑖_ updates are solved separately for each genomic block, while global **B** and **C** updates aggregate information across blocks. **B** and **C** are constrained to be non-negative to support interpretable positive loadings. Implementation details, including Khatri-Rao products, conjugate-gradient updates, SciPy LinearOperator use [40], ridge updates, and the non-negativity warm-up heuristic, are provided in Supplementary Method 4.

Operationally, **A**_𝑖_ captures which loci within block 𝑖 contribute to each latent axis, **B** captures the tissue profile of each axis, and **C** scales each axis in the phenotype mode. In the DR application, the columns of **B** define the dominant tissue anchoring of each inferred axis, while **A**_𝑖_ provides the gene and variant prioritization used for downstream single-cell projection and molecular follow-up. The block-wise formulation is essential for genome-wide scalability because it avoids fitting a single dense variant-by-variant structure across the entire genome.

The graph Laplacian term penalizes abrupt changes in **A**_𝑖_ across variants connected in the LD graph, while ridge penalties on **B** and **C** stabilize global factor estimates. Non-negativity supports axis weights that can be read as positive contributions rather than arbitrary signed rotations. These choices do not guarantee a global optimum, but they reduce rotational ambiguity and produce stable coordinate systems for downstream biological interpretation.

### Rank selection and solution stability

For standard CP decomposition, the Kruskal condition [42],

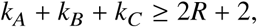

guarantees uniqueness under appropriate factor k-ranks. In the present single-phenotype setting, **C** ∈ ℝ^1×𝑅^ is a scaling vector, so 𝑘_𝐶_ = 1 and the Kruskal condition degenerates for 𝑅 ≥ 2. PRISMA therefore promotes practical identifiability through three complementary constraints: non-negativity, LD graph regularization, and empirical solution stability across random initializations. This should be interpreted as practical stabilization rather than a formal global-identifiability proof.

Rank selection used the Core Consistency Diagnostic (CORCONDIA) of Bro and Kiers [45], interpreted cautiously because the current DR application has a singleton phenotype mode. In compact form, CORCONDIA compares the estimated optimal core tensor **G**_𝑜𝑝𝑡_ with the superdiagonal reference core **G**_𝑠𝑢𝑝𝑒𝑟_:

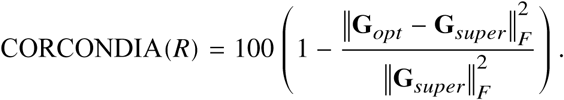

Ranks 𝑅 = 1 to 5 were evaluated by jointly considering CORCONDIA, variance explained, parsimony, and biological interpretability. Variance explained was defined as:

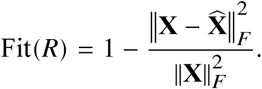

Rank 3 was selected as the lowest rank satisfying the CORCONDIA criterion (88.1% > 80%) while explaining 85.0% of variance and yielding interpretable tissue-biased axes supported by independent single-cell transcriptomic analyses. Solution stability was evaluated using 50 random initializations and sensitivity to the graph Laplacian penalty 𝜆; axis identities remained stable across the evaluated regularization range. Full CORCONDIA derivation, random-initialization alignment, cosine-similarity calculations, and lambda-sensitivity results are provided in Supplementary Method 5.

The final rank was not chosen by maximizing variance explained alone. Higher-rank solutions reconstructed more variance but were treated as candidate overfitting regimes unless supported by additional biological structure. Rank 3 was therefore selected as a parsimonious solution that crossed the model-consistency threshold, captured the major variance-explained elbow, and produced interpretable tissue-specific axes. This rank-selection rule balances mathematical fit and biological readability, which is particularly important in the single-phenotype setting where conventional CP uniqueness diagnostics are only partially applicable.

Empirical stability was assessed after optimal permutation alignment of factors across runs. The 50-initialization analysis evaluated whether axes reappeared with the same tissue identities and loading profiles despite the non-convex loss surface. The lambda-sensitivity analysis evaluated whether inferred axes depended strongly on graph regularization strength. Together, these checks support solution stability for the DR application, while avoiding claims of formal identifiability or global optimality.

### Simulation studies

Simulation studies evaluated null calibration, statistical power, SNP recovery, and runtime behavior under realistic matched GWAS/eQTL Z-score inputs. Synthetic GWAS and eQTL Z-scores were generated for 2,000 SNPs across 4 tissues, then combined into PRISMA integration scores. Under the null, independent simulations estimated the global loading distribution and empirical false-positive rates at nominal 𝛼 ∈ {0.01, 0.05, 0.10}. For power analyses, two orthogonal tissue-specific trajectories were injected at varying signal strengths, and power was defined as the fraction of causal SNP entries exceeding the standardized loading threshold. ROC/SNP recovery and runtime benchmarking were evaluated using the same factor-loading framework. Simulation outcomes are reported in Fig. 2 and Supplementary Note 1; full simulation design, thresholding, and runtime benchmark details are provided in Supplementary Method 6.

The simulation framework separates null calibration from signal recovery. Null datasets were used to estimate the distribution of standardized factor loadings and assess Type I error control, whereas independent signal-injected datasets were used for power and SNP-recovery analyses. This separation avoids circular use of the same simulations for both calibration and evaluation. Runtime benchmarks used real-data-scale tensor inputs to evaluate whether the block-wise solver could process genome-wide summary-statistics data on standard workstation hardware.

### Single-cell transcriptomic analyses

To assess whether PRISMA-derived axes map to relevant cellular compartments, factor-loading gene sets were projected onto three independent scRNA-seq atlases (Supplementary Table 3): fibrovascular membrane (FVM) samples from proliferative diabetic retinopathy and proliferative vitreoretinopathy (GSE165784) [46], PBMCs from DR patients and controls (GSE248284) [47], and a human retinal atlas containing photoreceptors, bipolar cells, and other retinal neurons (GSE137537) [48]. These datasets were not used to fit PRISMA; they provide orthogonal biological support for the inferred axes.

For each PRISMA trajectory, the top 200 genes ranked by absolute factor loading were used as axis-specific signatures. Per-cell module scores were computed using scanpy.tl.score_genes [49], and module-score differences were evaluated across dataset-defined contrast groups within annotated cell types. Empirical gene-set permutation tests used a PRISMA candidate-gene background to ask whether observed cell-type enrichment exceeded that expected from random candidate gene sets of matched size. The retinal atlas contains age-related macular degeneration rather than DR samples; it was therefore used to evaluate retinal cell-type enrichment rather than DR-specific dysregulation. These analyses support biological interpretation but do not establish whether the genetic variants causally drive the observed expression patterns. Dominance filtering, exact permutation P-value definitions, multiple-testing considerations, and gene-set-size sensitivity analyses are provided in Supplementary Method 7.

The FVM and PBMC analyses used dataset-defined contrast groups, whereas the retinal atlas analysis focused on enrichment across retinal cell classes. This distinction is important because GSE165784 is a fibrovascular membrane dataset rather than a healthy-retina control comparison, and GSE137537 is not a DR-specific atlas. Consequently, the single-cell analyses are interpreted as orthogonal support that PRISMA gene sets localize to plausible cellular compartments, not as proof of disease causality or direct replication of bulk eQTL effects in diseased cells.

### Post-GWAS comparator analyses

To evaluate whether PRISMA-prioritized targets showed concordance with established post-GWAS gene-level evidence, we performed MAGMA, SMR/HEIDI, and targeted colocalization analyses. Because the 549 PRISMA target rows corresponded to 405 unique gene symbols, gene-level comparator analyses used this 405-gene projection rather than duplicated target rows. MAGMA was used to assess GWAS gene-level and gene-set burden for the PRISMA target set, the three axis-specific gene sets, and an exploratory subthreshold target set. Genes not mappable to MAGMA Entrez identifiers were excluded from MAGMA gene-set testing and reported as a mapping limitation. SMR/HEIDI was used as a formal GWAS-eQTL gene-level comparator, with strict support defined as all-tissue BH-adjusted SMR q < 0.05 and HEIDI P > 0.01. Colocalization was not performed genome-wide; targeted coloc.abf analyses were restricted to QC-passed gene-tissue pairs among strict SMR-supported PRISMA genes to assess local shared-variant support. Colocalization support was defined as PP.H4 ≥ 0.80, with 0.50 ≤ PP.H4 < 0.80 treated as suggestive. Tissue-source caveats and sensitivity analyses are described in the Supplementary Methods and Supplementary Tables 24-28. These analyses were interpreted as post-GWAS concordance and sensitivity analyses, not as causal validation of all PRISMA targets.

### Benchmarking and height GWAS negative control

PRISMA was benchmarked against PCA, NMF [41], and K-means, with baseline implementations run using scikit-learn [52], applied to the same PRISMA integration-score matrix of 8,201 SNPs across 5 tissues. All methods were configured to extract 3 components. Tissue specificity was summarized using the Gini coefficient. For sorted tissue weights 𝑤_1_, 𝑤_2_, …, 𝑤_𝑇_, the Gini coefficient is:

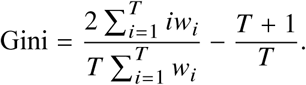

Mean Gini coefficients across components were used to compare tissue-regulatory resolution across methods. Baseline preprocessing, initialization settings, and scRNA-seq enrichment benchmarking details are provided in Supplementary Method 8.

The benchmarking analysis compared tissue-resolution properties under matched input data and rank, not whether PCA, NMF, or K-means solve the same biological problem as PRISMA. PCA provides orthogonal variance components, NMF provides non-negative matrix factors, and K-means provides cluster centroids; none incorporates LD graph topology directly. PRISMA was therefore evaluated in terms of tissue specificity and downstream biological enrichment rather than generic reconstruction error alone.

As a negative-control trait analysis, the identical PRISMA pipeline was applied to adult height GWAS summary statistics from GWAS Catalog accession ebi-a-GCST90018959 [12,13], corresponding to European ancestry and N=360,388 [12]. Height was analyzed using the same five-tissue eQTL panel as DR. The height tensor was decomposed with R=3, and height axes were matched to DR axes by dominant tissue loading. We compared top-200 gene overlap and rank shifts between matched DR and height axes to evaluate whether PRISMA captures trait-dependent regulatory reprioritization within a shared eQTL-mappable tissue framework. Jaccard, hypergeometric overlap, and rank-shift procedures are described in Supplementary Method 9.

This negative-control analysis asks whether the same eQTL-mappable tissue framework produces identical gene prioritization across unrelated polygenic traits. Substantial overlap is expected because both analyses draw from the same regulatory universe, but trait-dependent rank shifts indicate that GWAS signals reprioritize genes within that shared tissue framework. The analysis therefore supports regulatory reprioritization rather than binary tissue specificity.

### Exploratory vitreous multi-omics analysis

Exploratory vitreous proteomic and metabolomic analyses tested whether PRISMA-defined axes nominated molecular correlates in the disease microenvironment. The cohort comprised 13 vitrectomy samples: 7 from patients with severe diabetic retinopathy and 6 from non-diabetic controls. Targeted proteomics quantified 429 proteins whose encoding genes overlapped the union of PRISMA factor-loading gene sets. Proteins differing between DR and control samples were mapped back to their corresponding PRISMA trajectories, and per-sample proteomic module scores were computed by averaging row-wise Z-scores of significant proteins within each trajectory’s gene set.

Untargeted metabolomic analyses were restricted to the 7 DR samples for within-disease heterogeneity. For each retained metabolite feature, Spearman correlations were computed against the three trajectory module scores. Benjamini-Hochberg FDR correction [50] was applied by trajectory axis using statsmodels [51], and leave-one-out cross-validation was used as an exploratory sensitivity screen. These analyses were hypothesis-generating: the small DR metabolomics sample size (n=7) does not establish causal directionality, stable biomarkers, or clinical utility. Detailed compound annotation filtering, FDR procedures, trajectory specificity score *S*_𝑚_, LOOCV criteria, and final sensitivity-filtering rules are provided in Supplementary Method 10.

Proteomic analyses used all 13 samples for DR-versus-control module-score comparisons, whereas metabolomic correlation analyses were restricted to DR samples to focus on axis-associated heterogeneity among affected individuals. Annotated metabolite identities were treated as putative features because untargeted metabolomics can produce ambiguous compound assignments. The resulting proteomic and metabolomic findings are therefore presented as candidate molecular correlates of PRISMA axes and as motivation for future validation in larger, longitudinally sampled cohorts.

## Data Availability Statement

All GWAS summary statistics and GTEx eQTL data used in this study are publicly available. The FVM (GSE165784), PBMC (GSE248284), and retina (GSE137537) single-cell RNA-sequencing datasets are accessible through the Gene Expression Omnibus. Processed PRISMA weight matrices, cell-type scoring results, derived supplementary data, and the author-generated FinnGen+MVP diabetic retinopathy GWAS meta-analysis summary statistics are available at Zenodo (version-specific DOI: https://doi.org/10.5281/zenodo.20340998; all-versions DOI: https://doi.org/10.5281/zenodo.20340997).

## Code Availability Statement

The complete PRISMA computational framework, including the graph Laplacian-regularized factorization solver and downstream evaluation scripts, is implemented in Python. Source code, documentation, and a reproducible tutorial are available at https://github.com/DrHaoXiong/PRISMA.

## Abbreviation

CNS: Central Nervous System
DR: Diabetic Retinopathy
eQTL: expression quantitative trait locus
GWAS: genome-wide association study
INFO: Imputation Quality Score
IVW: inverse-variance weighted
LD: Linkage Disequilibrium
MHC: Major Histocompatibility Complex
MVP: Million Veteran Program
NVU: Neurovascular Unit
SMR: Summary-data-based Mendelian Randomization
SNP: single-nucleotide polymorphism
SuSiE: Sum of Single Effects

## Supporting information

supplementary table summary

## Acknowledgments

We thank the investigators and participants of FinnGen, the Million Veteran Program, GTEx, GEO, and the GWAS Catalog resources used in this study. We thank Mr. Yijing Liu for assistance with the computational hardware used for model testing.

## Author Contributions

H.X. conceived and designed the study and drafted the original manuscript. L.Z. and Z.W. curated the data. W.X. and A.J. prepared the figures and visualizations. H.X. contributed to the methodology, software development, and formal analysis. S.L., Z.X. and L.Z. performed data validation. J.Y. and Z.W. supervised the study. H.X. was responsible for project administration. All authors reviewed the manuscript and approved the final version for publication.

## Consent for publication

Not applicable.

