## Supplementary material for "PRISMA: A tensor-based framework for deconstructing the genetic architecture of complex diseases, with application to diabetic retinopathy": Supplementary_Data_Legends.txt

# Supplementary Data Legends

## Supplementary Data 1 | Full PRISMA Factor A SNP loadings

Machine-readable complete locus-mode factor loading matrix from the final PRISMA decomposition. Rows correspond to SNP-level instruments retained after GWAS-eQTL integration, gene-level SMR pruning, and targeted purification. Columns report Factor A loadings for Rank 0, Rank 1, and Rank 2, corresponding to the three PRISMA-derived genetic axes used throughout the manuscript.

---

## Supplementary Data 2 | Figure 7 metabolomics source data

Machine-readable source data exported from the Figure 7 metabolomics plotting script. Rows are keyed by panel and include the full volcano-plot correlation data, specificity-versus-LOOCV scatter data, and top FDR-ranked metabolite-axis associations used for the bar plot. This file documents the plotting values underlying Fig. 7 and links the figure to Supplementary Tables 9 and 11.

---

## Supplementary Data 3 | Figure 2 simulation source data

Machine-readable source data exported from the April 18 enhanced Figure 2 simulation code path. Rows are keyed by panel and include Type I error calibration, statistical power, ROC coordinates with the corresponding ROC AUC, and computational-efficiency measurements. The fixed rerun seed is recorded to make the frozen Figure 2 benchmark auditable and reproducible.

---

## Supplementary Data 4 | Figure 5 single-cell source data

Machine-readable cell-level source data for the real-score Figure 5 single-cell spatial mapping. Rows include UMAP coordinates, cell type, condition where available, PRISMA module score, dataset, axis label, and the exact score-source file used for FVM Axis 1, PBMC Axis 2, and retinal Axis 3 panels. This file replaces legacy mock/fallback UMAP score tables and links Figure 5 directly to the cell-level module-score outputs underlying Supplementary Tables 8 and 15.
