## supplementary figures for "PRISMA: A tensor-based framework for deconstructing the genetic architecture of complex diseases, with application to diabetic retinopathy": Supplementary_Figure_1_Baseline_Comparison.pdf

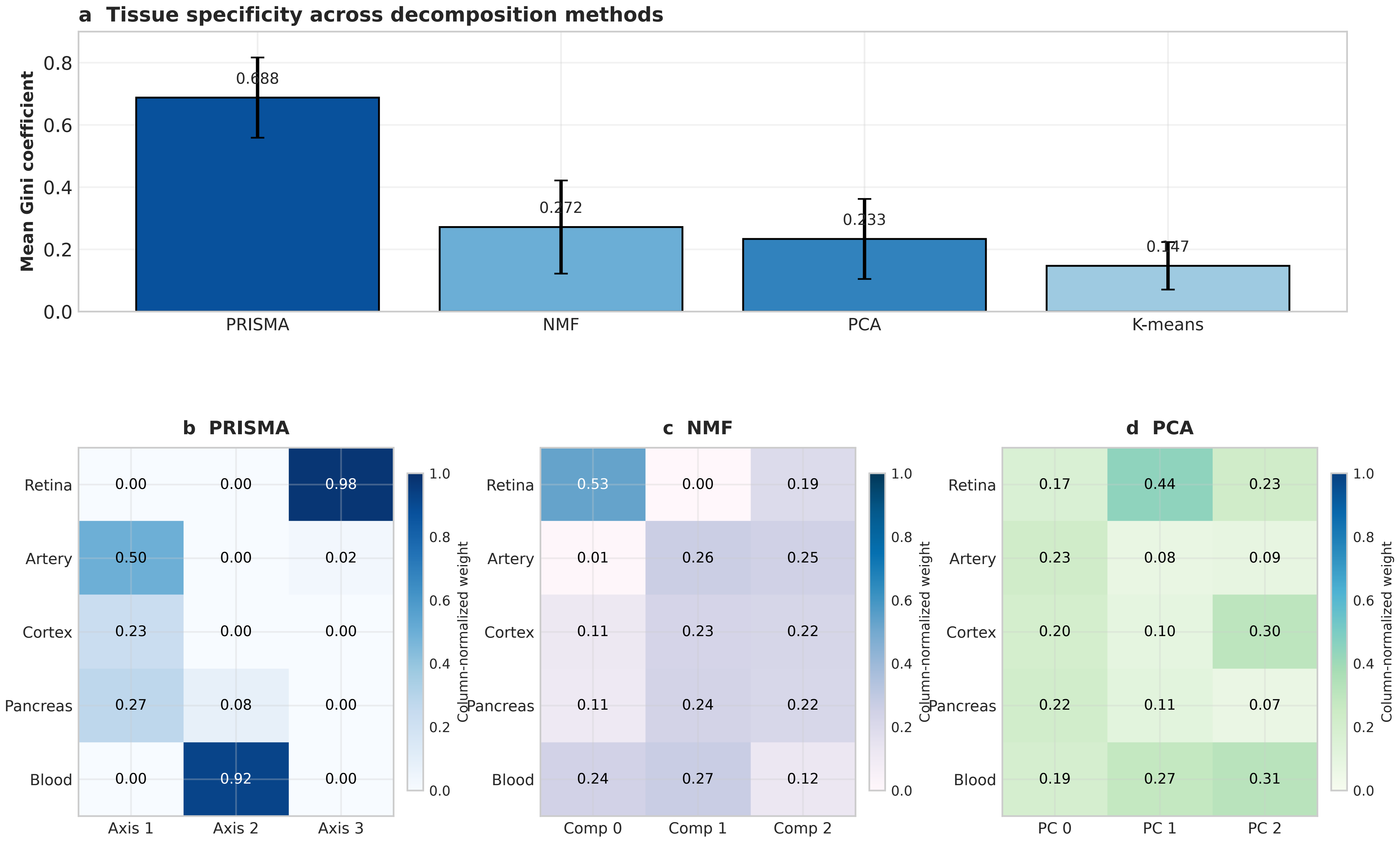


PRISMA rank order follows the current manuscript and Supplementary Table 12: Axis 1 = vascular-metabolic, Axis 2 = immune-inflammatory, Axis 3 = retina-specific neurodegenerative.
