## supplementary figures for "PRISMA: A tensor-based framework for deconstructing the genetic architecture of complex diseases, with application to diabetic retinopathy": Supplementary_Figure_Legends.txt

# Supplementary Figure Legends

## Supplementary Figure 1 | Baseline method comparison demonstrates PRISMA's superior tissue-regulatory resolution.

**a**, Gini coefficient comparison across four methods (PRISMA, NMF, PCA, K-means) quantifying tissue specificity of extracted components. PRISMA achieves substantially higher tissue specificity (Gini = 0.688 ± 0.129) compared to NMF (0.272 ± 0.150), PCA (0.233 ± 0.129), and K-means (0.147 ± 0.076), demonstrating 2.5-fold to 4.7-fold improvement in tissue-anchored decomposition. Higher Gini coefficients indicate stronger concentration of factor weights in specific tissues rather than diffuse multi-tissue patterns.

**b-d**, Tissue weight heatmaps for PRISMA, NMF, and PCA. **b**, PRISMA produces sharp, interpretable tissue-specific loadings (e.g., Axis 2: whole blood weight = 0.996; Axis 3: retina weight = 0.999). **c**, NMF shows moderate tissue clustering but lacks clear dominant tissues per component. **d**, PCA produces diffuse, multi-tissue patterns with no interpretable tissue specificity. K-means is excluded from heatmap visualization as it performs clustering rather than factor decomposition, producing discrete cluster assignments rather than continuous tissue weights; its Gini coefficient (0.147 ± 0.076) is included in panel **a** for quantitative comparison.

The superior tissue specificity of PRISMA stems from graph Laplacian regularization, which explicitly preserves local linkage disequilibrium (LD) structure—a biological prior entirely absent in conventional dimensionality reduction methods. By enforcing that SNPs in high LD share coordinated factor loadings, PRISMA captures genuine regulatory co-inheritance patterns rather than arbitrary statistical variance. This demonstrates that the observed tissue-anchored decomposition is not an artifact of simple clustering algorithms or driven merely by differences in eQTL sample sizes across tissues, but reflects biologically meaningful regulatory architectures.

---

## Supplementary Figure 2 | Height GWAS negative control demonstrates disease-specific target prioritization.

To empirically test whether PRISMA's decomposition merely captures generic tissue expression architectures rather than genuine disease-driven etiology, we applied the identical tensor framework to adult height GWAS summary statistics (GIANT consortium, N=360,388) using the same five-tissue eQTL panel. Height was selected as a negative control because it represents a highly polygenic anthropometric trait with fundamentally distinct genetic architecture from diabetic retinopathy—height variants primarily influence skeletal development and growth hormone pathways, whereas DR variants target vascular, immune, and neural pathways.

**a**, Tissue specificity comparison between DR and Height Factor B weights in the matched negative-control rerun. Both traits achieve comparable tissue-anchored decomposition (Height: CORCONDIA = 64.6%, mean Gini = 0.687; matched DR rerun: CORCONDIA = 64.6%, mean Gini = 0.688), confirming mathematical generalizability without conflating this rerun with the primary DR rank-selection analysis. However, the dominant tissues differ between traits: DR Axis 1 is anchored in artery and pancreas (vascular-metabolic), while Height shows distinct tissue prioritization patterns reflecting skeletal and growth-related regulatory networks.

**b**, Gene overlap analysis across matched axes. The overlap is substantial (39-45%) relative to the shared eQTL-mappable gene universe (N = 8,177; hypergeometric test, all P < 10^-77), reflecting a shared tissue-regulatory backbone common to polygenic traits that utilize overlapping eQTL architectures rather than disease-specific equivalence. Shared genes (dark blue) represent the common regulatory infrastructure, while trait-specific genes (light blue: DR only; very light blue: Height only) demonstrate divergent pathogenic routes within the same tissue framework.

**c**, Rank-shift scatter plot demonstrating trait-specific prioritization. Each point represents a gene shared between DR and Height, with coordinates showing its rank in each trait (1 = highest weight). Deviation from the diagonal (dashed line = perfect rank consistency) indicates trait-specific reprioritization. PRISMA's trait-specificity manifests not in absolute gene presence or absence, but in relative weight prioritization: genes receiving high weights in DR (e.g., TAGLN, ALDH16A1) are present in Height's gene pool but rank lower, while Height-prioritized examples (e.g., GBP3, CUTALP) show the inverse pattern. Representative rank-shift records are provided in Supplementary Table 19. This rank-shift demonstrates that PRISMA dynamically reprioritizes regulatory networks based on trait-specific GWAS signals, rather than merely clustering ubiquitous eQTL effects.

Collectively, these results confirm that the mathematical weights output by PRISMA reflect genuine, trait-specific pathogenic leverage within a shared tissue-regulatory framework. The successful decomposition of both DR and Height, combined with clear trait-specific reprioritization, provides proof-of-principle for trajectory-resolved genetic dissection across diverse polygenic traits.

---

## Supplementary Figure 3 | Permutation test sensitivity analysis confirms robustness of spatial enrichment.

To ensure that the spatial localization of PRISMA Axis 1 (vascular-metabolic) in fibrovascular membrane (FVM) pathological cells was not an artifact of arbitrary gene-set size thresholding, we performed a rigorous permutation stress-test across sliding cutoffs (Top 50, 100, 150, 200, and all 243 strictly Axis 1-dominant genes). For each cutoff, we computed module scores using scanpy.tl.score_genes and compared the observed enrichment (FVM pathological cells vs. other cells) against 1,000 random gene sets of equal size sampled from the disease-associated background.

**a-e**, Empirical null distributions (gray histograms) of mean difference in module scores (Δ Mean) for each gene-set size. Red dashed vertical lines indicate the observed effect size using PRISMA Axis 1 genes, with corresponding empirical P-values. The pathological spatial enrichment remains robust across all tested thresholds (P = 0.010; Supplementary Table 16), confirming that PRISMA's topological weights capture genuine biological signal rather than statistical noise.

**f**, Summary bar plot showing -log₁₀(empirical P-value) across all tested cutoffs. All tested cutoffs meet the prespecified robustness threshold (P ≤ 0.01), demonstrating that the spatial localization is stable and not dependent on arbitrary parameter choices. This robustness validates that the graph Laplacian regularization successfully anchors the latent factors to biologically coherent, tissue-specific regulatory programs.
