## Supplementary figures and images for "PRISMA: A tensor-based framework for deconstructing the genetic architecture of complex diseases, with application to diabetic retinopathy"

### Supplementary_Figure_1_Baseline_Comparison.png

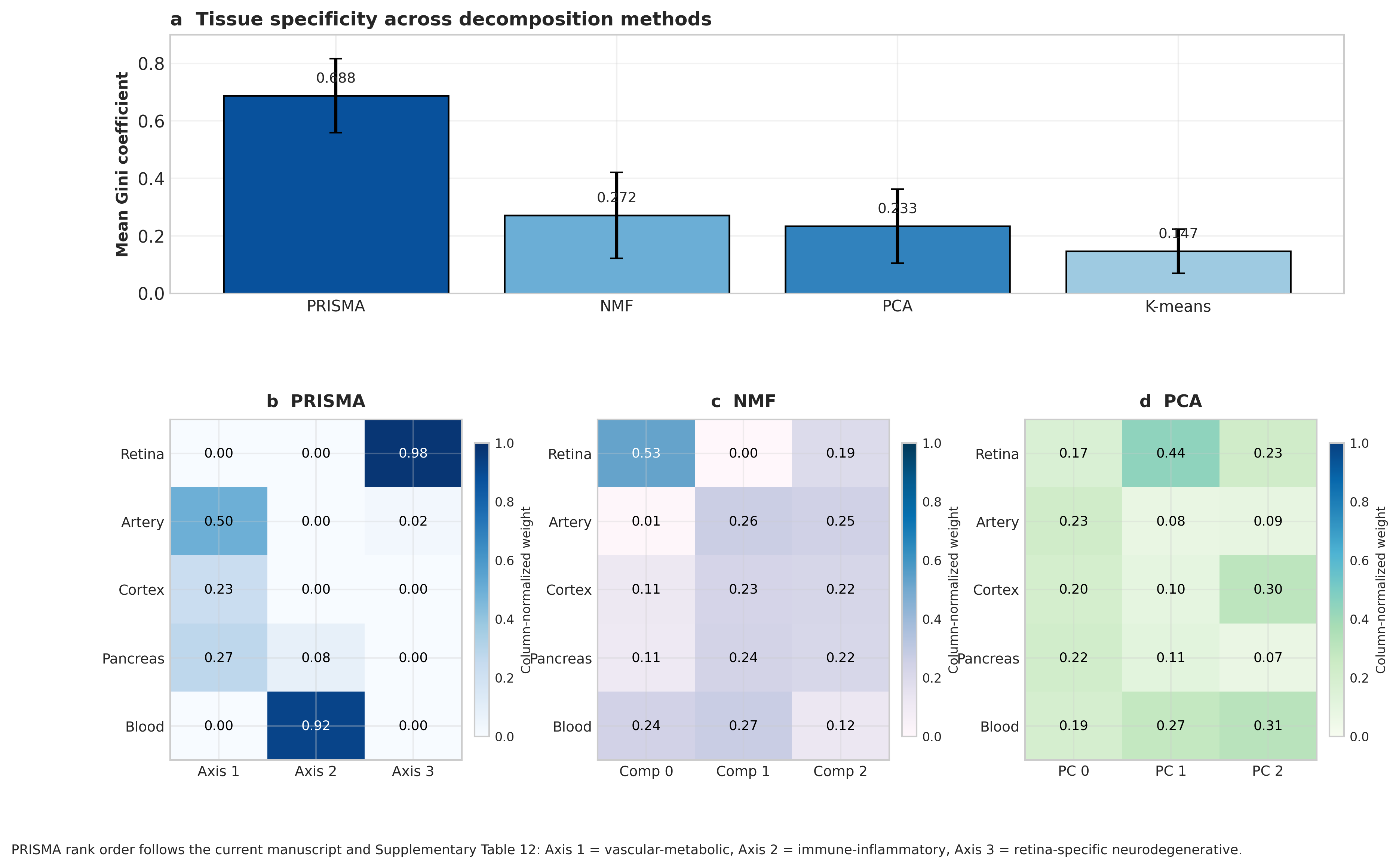

### Supplementary_Figure_2_Height_Negative_Control.pdf

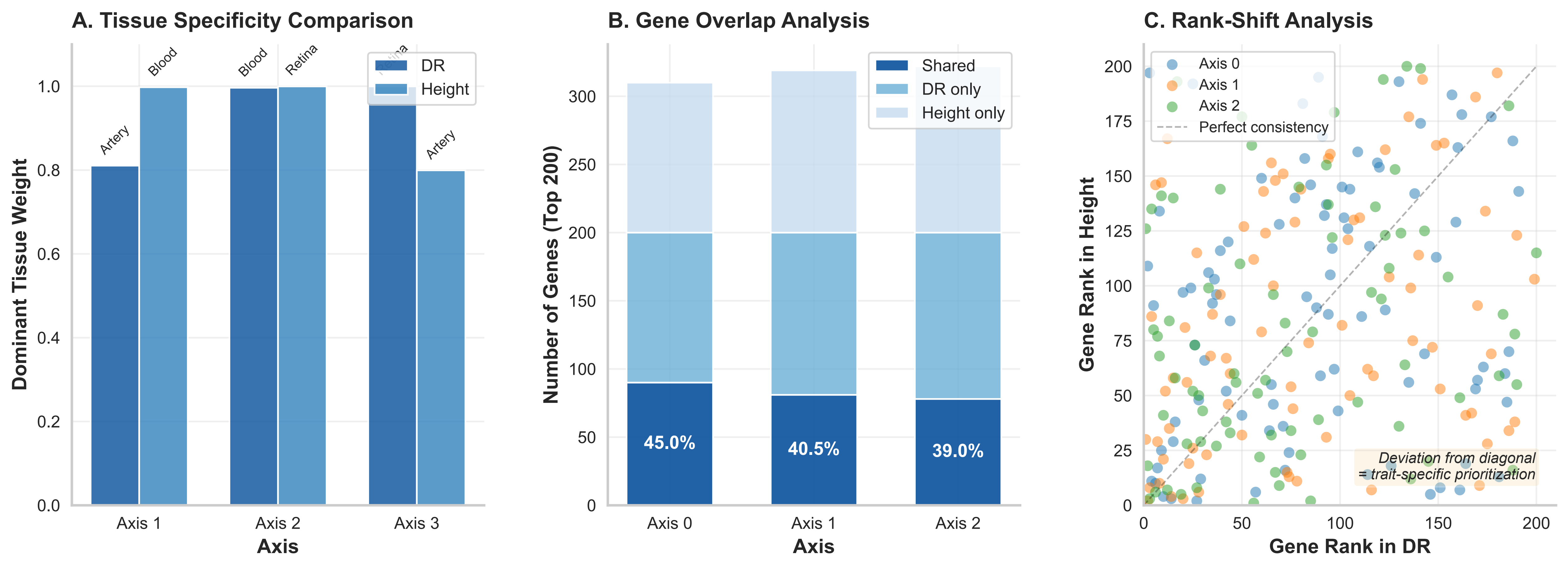

### Supplementary_Figure_2_Height_Negative_Control.png

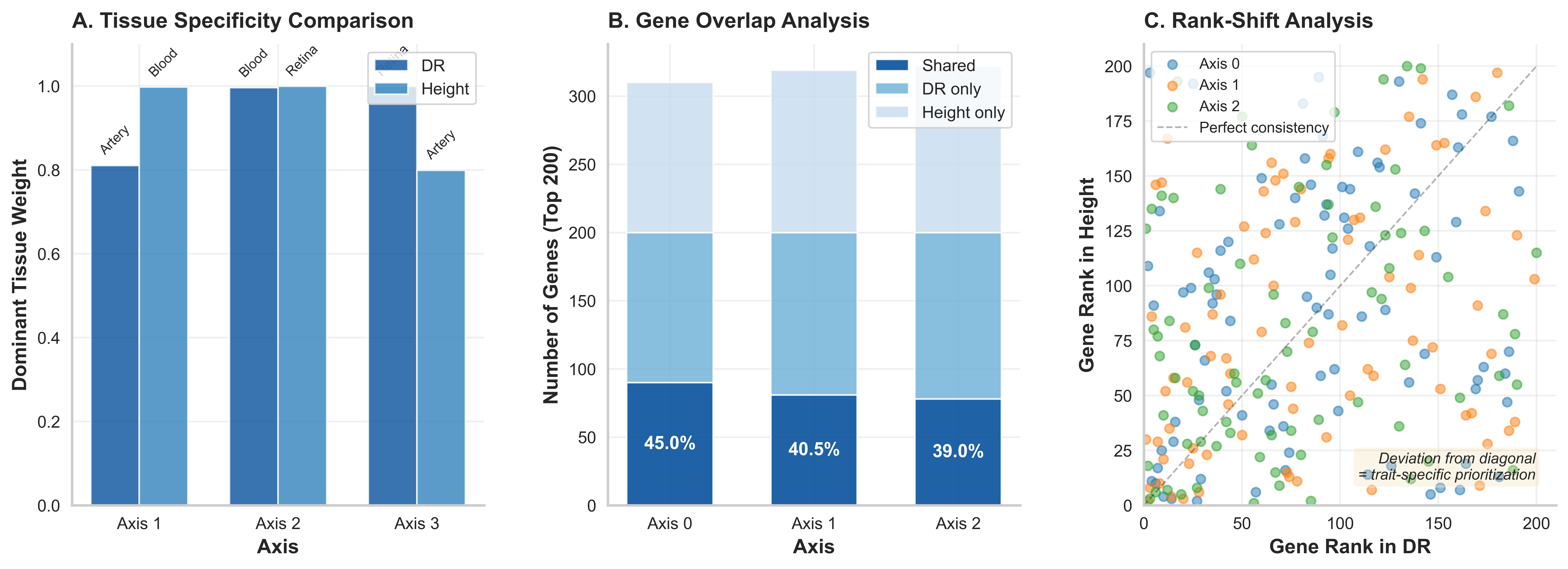

### Supplementary_Figure_3_Permutation_Sensitivity.pdf

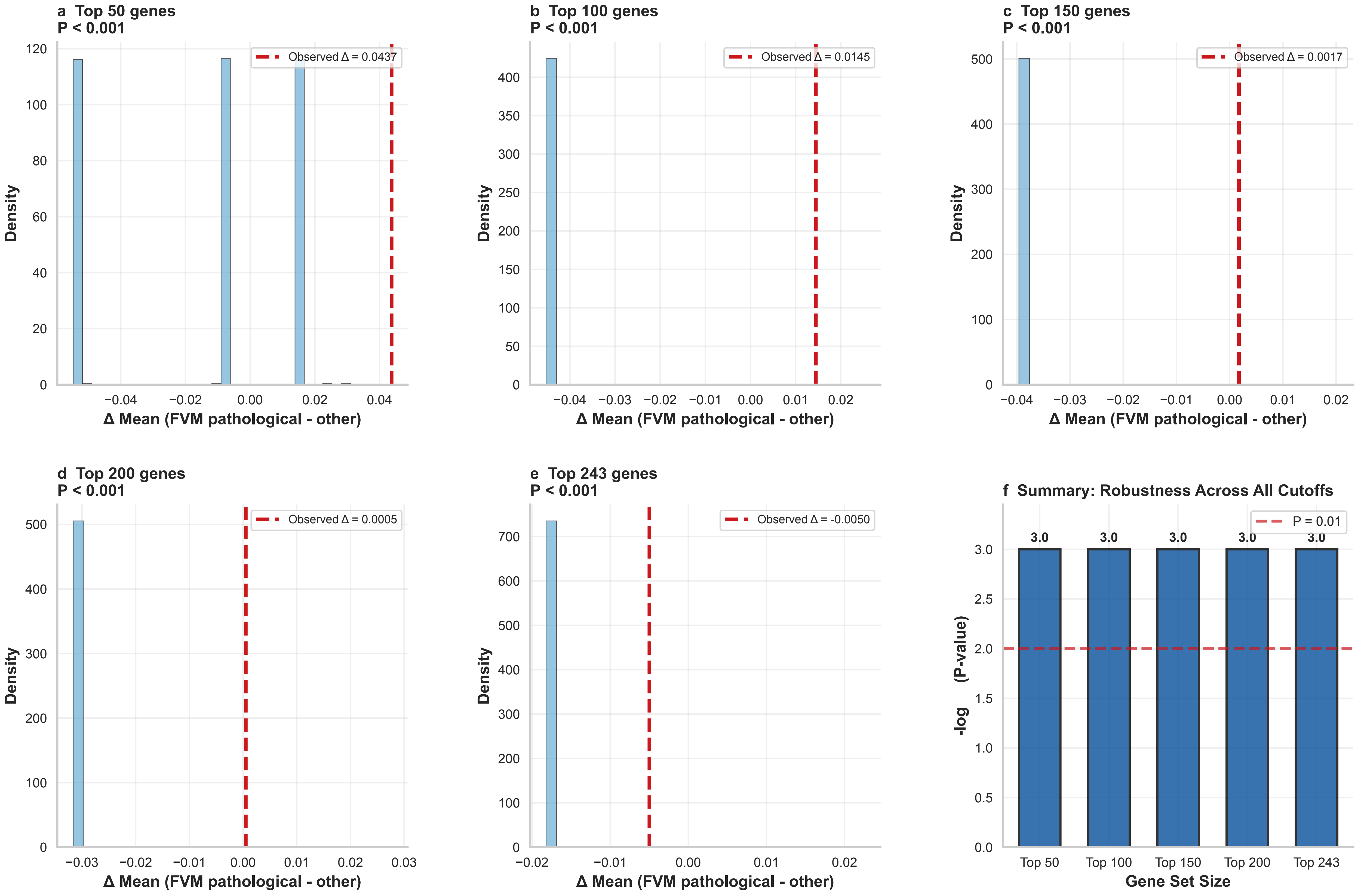

### Supplementary_Figure_3_Permutation_Sensitivity.png

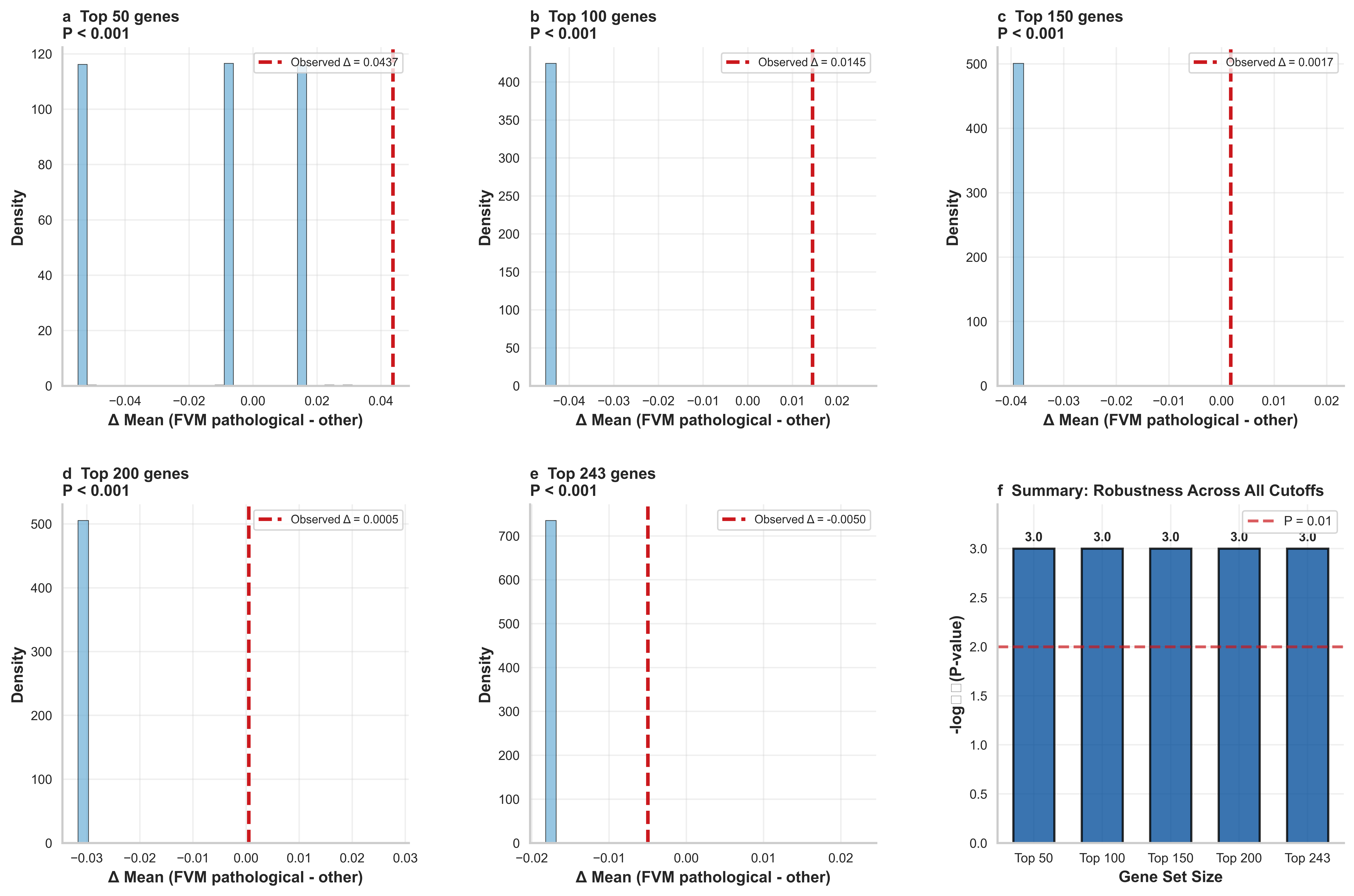
