## supplementary methods for "PRISMA: A tensor-based framework for deconstructing the genetic architecture of complex diseases, with application to diabetic retinopathy"

#### Supplementary Method 1. GWAS and eQTL harmonization

Diabetic retinopathy (DR) GWAS summary statistics were obtained from two European-ancestry sources: the FinnGen research project (R12) [30] and the Million Veteran Program (MVP) [31,32] (Supplementary Table 1). A fixed-effect inverse-variance weighted meta-analysis was performed using METAL [33], yielding a combined sample size of $N = 505,435$. Standard GWAS quality-control filters removed variants with minor allele frequency < 0.01, imputation quality (INFO) < 0.9, or standard error $\leq 0$. Strand-ambiguous variants and variants in the major histocompatibility complex (MHC; chr6:25-35 Mb) were excluded. Genomic coordinates were referenced to GRCh37, and alleles were harmonized to the 1000 Genomes Project Phase 3 European reference panel [34].

Cis-eQTL summary statistics for five DR-relevant tissues were used to construct the PRISMA tensor (Supplementary Table 2). Retina-specific eQTLs were obtained from a human retina atlas [35]; eQTLs for tibial artery, brain cortex, pancreas, and whole blood were obtained from GTEx Release v8 [36]. For each tissue, eQTL Z-scores were computed from reported effect sizes and standard errors. If the genomic inflation factor $\lambda_{GC}$ exceeded 1.0, eQTL Z-scores were rescaled by $1/\sqrt{\lambda_{GC}}$ before integration.

Alleles were aligned to the GWAS backbone using the SMR allele-matching protocol [2]. Effect alleles (A1) and reference alleles (A2) were harmonized between GWAS and eQTL panels. For variants with flipped alleles, the signs of corresponding eQTL effect sizes and eQTL summary statistics were inverted to maintain directional consistency. For each gene, only the strongest cis-eQTL, defined by cross-tissue composite $|Z|$, was retained as the sentinel instrumental variable. This gene-level sentinel selection minimizes collinearity within local genomic windows and provides a compact regulatory input for PRISMA feature construction.

To reduce broadly expressed or structurally confounded signals, genes encoding ribosomal proteins (RPS*, RPL*) and translation elongation/initiation factors (EEF*, EIF*) were excluded as a pre-specified filter for translation-related housekeeping signals [37]. Genes in the 17q21.31 inversion region (*KANSL1-AS1, ARHGAP27, LRRC37A2, ARL17A, MAPT, CRHR1, SPPL2C, STH*) were also excluded to mitigate long-range LD artifacts [38]. The genome was partitioned into approximately 1,363 independent LD blocks defined by LDetect for the European population in hg19 coordinates [39]. Blocks overlapping complex LD regions, including MHC chr6:25-35 Mb, 8p23 inversion chr8:7-13 Mb, and 11p15.5 imprinting domain chr11:45-57 Mb, were excluded from tensor construction.

#### Supplementary Method 2. PRISMA GWAS-eQTL integration score

PRISMA maps locus-level disease associations to tissue-specific regulatory contexts by integrating GWAS and eQTL summary statistics using a PRISMA-specific bounded Z-score formula. After allele harmonization, each variant-gene-tissue combination receives a standardized GWAS-eQTL integration score, denoted $S_{int}$:

$$S_{int} = \mathrm{sign}(Z_{GWAS} Z_{eQTL}) \sqrt{\frac{Z_{GWAS}^2 Z_{eQTL}^2}{Z_{GWAS}^2 + Z_{eQTL}^2 + \varepsilon}}$$

Here, $Z_{GWAS}$ is the disease association Z-score, $Z_{eQTL}$ is the tissue-specific eQTL Z-score, and $\varepsilon = 10^{-8}$ is included for numerical stability when both Z-scores are near zero. The sign convention is $\mathrm{sign}(0) = 0$. Variants with aligned disease and regulatory directions receive positive scores, variants with opposing directions receive negative scores, and variants weakly supported by either GWAS or eQTL evidence receive attenuated scores because of the harmonic-mean-like denominator.

The score is used as a feature weight for tensor decomposition. It is not interpreted as formal causal SMR evidence. Formal causal SMR analysis would require additional heterogeneity testing, such as HEIDI, and stronger assumptions about instrument validity. In PRISMA, this algebraic form is used to prioritize variant-gene-tissue entries jointly supported by disease association and local expression regulation before downstream factorization. Sub-threshold targets prioritized by this score and subsequent factorization are candidate signals for tissue-aware follow-up, not newly discovered genome-wide significant loci.

#### Supplementary Method 3. Tensor construction and LD graph construction

Following targeted purification, one-dimensional GWAS-eQTL integration scores are projected into block-wise multidimensional structures. For each genomic block $i$, PRISMA constructs a variant-by-tissue association matrix:

$$\mathbf{X}_i \in \mathbb{R}^{N_i \times T}$$

where $N_i$ is the number of variants in block $i$ and $T$ is the number of profiled tissues. PRISMA is specified for a general three-way tensor:

$$\mathcal{X} \in \mathbb{R}^{N \times T \times P}$$

where $N$ is the genetic-variant dimension, $T$ is the tissue dimension, and $P$ is the phenotype dimension. The present DR application evaluates the single-phenotype case (DR, $P = 1$). In this setting, the phenotype mode is a singleton dimension and the phenotype-mode factor reduces to a rank-specific scaling vector. Each genomic block is therefore represented operationally as $\mathbf{X}_i \in \mathbb{R}^{N_i \times T}$, while retaining the tensor notation for the general model specification. Multi-phenotype implementations, including sample-overlap correction across GWAS cohorts, are architectural extensions to be evaluated in future work.

To reduce the influence of extreme statistical outliers, a variance-stabilizing square-root transformation is applied:

$$\tilde{\mathbf{X}}_i = \mathrm{sign}(\mathbf{X}_i)\sqrt{|\mathbf{X}_i|}$$

with transformed values bounded within $[-20, 20]$. This transformation preserves the continuous relative magnitude of moderate-effect SNPs while strictly bounding isolated mega-locus signals that could dominate the Frobenius norm during decomposition.

Local LD topology is encoded as a graph Laplacian. Using the 1000 Genomes Project Phase 3 European reference panel [34], PRISMA computes the Pearson correlation matrix $\mathbf{R}$ among variants within each block. The adjacency matrix is defined as:

$$\mathbf{W} = |\mathbf{R}|$$

with zero diagonal, and the corresponding graph Laplacian is:

$$\mathbf{L}_i = \mathbf{D}_i - \mathbf{W}_i$$

where $\mathbf{D}_i$ is the diagonal degree matrix. The absolute correlation matrix $|\mathbf{R}|$ is used rather than $\mathbf{R}^2$ so that arbitrary allelic phase is discarded while the linear magnitude of LD is preserved. This choice regularizes variants in strong LD, whether positively or negatively correlated, toward similar magnitudes of factor loading. Compared with $\mathbf{R}^2$, $|\mathbf{R}|$ avoids disproportionately shrinking intermediate LD values and preserves a more cohesive network topology for capturing coordinated regulatory effects.

#### Supplementary Method 4. Graph-regularized block-wise ALS implementation

PRISMA is implemented as a three-dimensional SNP × tissue × trait tensor framework. In the current single-trait DR application ($P = 1$), Factor C is estimated as a rank-specific scaling vector, making this instance algebraically equivalent to a graph-regularized coupled matrix factorization across genomic blocks. This equivalence reflects the single-trait instantiation of the general tensor model rather than a separate matrix-only algorithm.

For each genomic block $i$, PRISMA approximates the input tensor or block matrix $\mathbf{X}_i$ using low-rank latent factors:

$$\mathbf{A}_i \in \mathbb{R}^{N_i \times R}, \quad \mathbf{B} \in \mathbb{R}^{T \times R}, \quad \mathbf{C} \in \mathbb{R}^{1 \times R}$$

where $\mathbf{A}_i$ contains locus-level genetic drivers, $\mathbf{B}$ contains global tissue specificity weights, $\mathbf{C}$ contains global phenotypic relevance or rank-specific scaling, and $R$ is the predefined rank. The graph-regularized objective is:

$$\min_{\{\mathbf{A}_i\},\mathbf{B},\mathbf{C}} \sum_i \|\mathbf{X}_i - [[\mathbf{A}_i, \mathbf{B}, \mathbf{C}]]\|_F^2 + \lambda_{reg} \sum_i \mathrm{Tr}(\mathbf{A}_i^\top \mathbf{L}_i \mathbf{A}_i) + \lambda_b \|\mathbf{B}\|_F^2 + \lambda_c \|\mathbf{C}\|_F^2$$

For the single-phenotype case, $\mathbf{C}$ reduces to a $1 \times R$ scaling vector and the reconstruction simplifies to:

$$\mathbf{X}_i \approx \mathbf{A}_i \mathbf{D}_C \mathbf{B}^\top$$

where $\mathbf{D}_C = \mathbf{diag}(\mathbf{C})$. When $P > 1$, Factor C would become a $P \times R$ phenotype-mode matrix, but evaluation of that multi-phenotype setting is left to future work.

The objective is optimized using block-wise alternating least squares (ALS). Because the genetic dimension is much larger than the tissue dimension, $\mathbf{A}_i$ is updated locally for each LD block rather than globally across all variants. The local update uses the Khatri-Rao product:

$$\mathbf{K}_{BC} = \mathbf{B} \odot \mathbf{C}$$

and is solved with Krylov subspace methods, specifically conjugate gradient (CG) implemented through SciPy's LinearOperator [40]. This permits implicit matrix-vector multiplication without materializing dense $(N_i \times N_i)$ covariance matrices.

After local updates of $\mathbf{A}_i$, global matrices $\mathbf{B}$ and $\mathbf{C}$ are updated by aggregating sufficient statistics across all blocks and solving ridge-regularized least-squares subproblems. $\mathbf{B}$ and $\mathbf{C}$ are constrained to be non-negative to produce interpretable positive loadings. To reduce premature convergence to the boundary of the non-negative orthant, PRISMA uses a dynamic warm-up heuristic: the non-negativity constraint is delayed for the initial 30% of maximum epochs, allowing latent factors to traverse the loss landscape before projection onto the non-negative coordinate system. This relaxation-then-projection strategy is consistent with established non-convex optimization practice and with comparable strategies used in multiplicative updates for non-negative matrix factorization [41].

#### Supplementary Method 5. Rank selection and stability diagnostics

For standard CP decomposition, the Kruskal condition [42],

$$k_A + k_B + k_C \geq 2R + 2$$

where $k$ denotes the k-rank of each factor matrix, guarantees unique recovery of latent factors. In the current single-phenotype setting, $\mathbf{C} \in \mathbb{R}^{1\times R}$ serves as a scaling vector, so the model reduces to coupled matrix factorization and the Kruskal condition degenerates because $k_C = 1$ violates the inequality for

$R \geq 2$. PRISMA therefore relies on practical identifiability through non-negativity constraints, LD-informed graph Laplacian regularization, and empirical convergence stability.

Rank selection uses the Core Consistency Diagnostic (CORCONDIA) of Bro and Kiers [45]. CORCONDIA was originally designed for standard CP tensor decomposition; in the single-phenotype regime, where Factor C reduces to a scaling vector, it should be interpreted as a relative diagnostic of low-rank structure rather than a formal proof of tensor identifiability. Given factor matrices $\mathbf{A}, \mathbf{B}, \mathbf{C}$ estimated at rank $R$, the optimal core tensor is computed as:

$$\mathbf{G}_{opt} = \mathbf{A}^{+}\mathbf{X}_{(1)}(\mathbf{C} \odot \mathbf{B})^{+\top}$$

where $\mathbf{X}_{(1)}$ is the mode-1 unfolding of $\mathbf{X}$, $\mathbf{A}^{+}$ is the Moore-Penrose pseudoinverse of $\mathbf{A}$, and $\mathbf{C} \odot \mathbf{B}$ is the Khatri-Rao product. The reference tensor $\mathbf{G}_{super}$ is the superdiagonal identity tensor of order $R$. CORCONDIA is:

$$\text{CORCONDIA}(R) = 100 \times \left(1 - \frac{\|\mathbf{G}_{opt} - \mathbf{G}_{super}\|_F^2}{\|\mathbf{G}_{super}\|_F^2}\right)$$

Ranks $R \in \{1, \ldots, 5\}$ were evaluated by jointly considering CORCONDIA, variance explained, parsimony, and biological interpretability. CORCONDIA > 80% was used as a guideline for adequate model fit rather than an absolute threshold. For the DR dataset, rank 1 had CORCONDIA = 25.0% and explained 35.9% of variance, rank 2 had CORCONDIA = 79.3% and explained 74.6% of variance, and rank 3 satisfied the CORCONDIA criterion at 88.1% while explaining 85.0% of variance. Higher ranks explained more variance (R=4: CORCONDIA = 93.9%, variance = 92.3%; R=5: CORCONDIA = 99.2%, variance = 99.2%) but were treated as candidate overfitting regimes requiring biological justification beyond numerical reconstruction. The selected rank was therefore R=3. If no rank satisfies CORCONDIA > 80%, the fallback criterion is the elbow of the variance-explained curve:

$$\text{Fit}(R) = 1 - \frac{\|\mathbf{X} - \widehat{\mathbf{X}}\|_F^2}{\|\mathbf{X}\|_F^2}$$

with $\widehat{\mathbf{X}} = \mathbf{A}(\mathbf{C} \odot \mathbf{B})^{\top}$ reconstructed via the Khatri-Rao product.

Stability was assessed using 50 random initializations, sampled uniformly from $U(0, 1)$, on the DR dataset. For each pair of solutions, corresponding factor matrices were aligned by optimal permutation and compared using cosine similarity. Mean pairwise cosine similarities were 1.000 ± 7×10^-17 for Factor A, 1.000 ± 2×10^-17 for Factor B, and 1.000 ± 0 for Factor C (Supplementary Table 14).

Sensitivity to the Laplacian penalty weight $\lambda_{reg}$ was evaluated across $\{0.01, 0.1, 1, 10, 100\}$. Axis-to-tissue assignments were defined by $\arg\max_t B_{t,r}$ for each rank $r$, and the Adjusted Rand Index (ARI) was computed across settings. ARI remained 1.00 across the evaluated range. Dominant tissue weights showed limited variation: Axis 1 weights ranged from 0.810 to 0.824, Axis 2 from 0.959 to 0.996, and Axis 3 from 0.9997 to 1.0000 (Fig. 3c-e, Supplementary Tables 10 and 21). The production analyses used $\lambda_{reg} = 0.1$.

#### Supplementary Method 6. Simulation studies

Simulation experiments evaluated Type I error calibration, power, ROC/SNP recovery, and runtime behavior under realistic matched GWAS/eQTL Z-score inputs. Synthetic data were generated for 2,000 SNPs across 4 tissues. GWAS Z-scores were drawn as:

$$Z_p^{GWAS} \sim N(0,1)$$

and eQTL Z-scores as:

$$Z_{pt}^{eQTL} \sim N(0,1)$$

PRISMA integration scores were computed from these simulated GWAS and eQTL Z-scores. Signal was injected under a two-trajectory design. SNPs $p \in [1, 500]$ carried additive Gaussian perturbations of magnitude $s$ in tissues 1-2, and SNPs $p \in [501, 1,000]$ carried the same perturbation in tissues 3-4. These two injected groups represented orthogonal genetic trajectories with divergent tissue specificities; the remaining 1,000 SNPs were null across all tissues. The signal level $s$ took values in $\{0, 1.5, 2.5, 3.5, 4.5, 5.5\}$.

To establish the global null distribution, 50 independent null trials ($s = 0$) were run. The empirical mean $\mu_{global}$ and standard deviation $\sigma_{global}$ of the raw factor loading matrix $\mathbf{A}$ were estimated across all entries. This estimation set was kept separate from all subsequent test datasets. In Type I error trials, 50 additional null datasets were generated and standardized as:

$$\widetilde{A}_{pr} = (A_{pr} - \mu_{global})/\sigma_{global}$$

The empirical false positive rate at nominal level $\alpha$ was computed as the fraction of all $2,000 \times 2$ standardized entries satisfying $|\widetilde{A}_{pr}| > z_{\alpha/2}$, evaluated at $\alpha \in \{0.01, 0.05, 0.10\}$. For power assessment, 20 trials were run at each nonzero signal level, and power was defined as the fraction of the 1,000 injected signal SNP entries per component satisfying $|\widetilde{A}_{pr}| > 1.96$.

ROC/SNP recovery analyses treated injected signal SNP entries as positives and null SNP entries as negatives. Standardized factor-loading scores were used to evaluate discriminative performance across thresholds. Runtime benchmarking used the full real-data PRISMA setting described in the main manuscript, including 993,226 pre-LD candidate variants across five eQTL tissues, corresponding to 4,966,130 variant-by-tissue cells, and five full real-data runs on an Intel Core i9 14900K processor.

### Supplementary Method 7. Single-cell transcriptomic validation

Independent scRNA-seq atlases were used to assess whether PRISMA-derived genetic trajectories mapped to disease-relevant cell types (Supplementary Table 3). These datasets were independent of PRISMA model fitting: factor loadings were derived only from summary-level GWAS statistics and bulk-tissue eQTL data. The orthogonal analysis datasets were: a fibrovascular membrane (FVM) atlas derived from proliferative diabetic retinopathy and proliferative vitreoretinopathy fibrovascular membranes (GSE165784) [46], a PBMC atlas from DR patients and controls (GSE248284) [47], and a human retinal atlas containing photoreceptors, bipolar cells, and other retinal neurons (GSE137537) [48].

For each trajectory $r \in \{0, 1, 2\}$, the top 200 genes ranked by absolute factor loading in $\mathbf{A}$ were selected as the axis-specific signature. For Axis 3 analysis in the retinal atlas, a dominance filter was used to select genes with $|A_{p,2}| > |A_{p,0}|$ and $|A_{p,2}| > |A_{p,1}|$, yielding 796 Axis 3 dominant genes. Per-cell module scores were computed using `scanpy.tl.score_genes` (v1.9) [49]. This procedure calculates the mean expression of signature genes minus the mean expression of a size-matched control gene set, with $n_{ctrl} = 50$ control genes drawn from equivalent expression bins and applied to raw counts.

Module scores were compared between dataset-defined contrast groups using the Wilcoxon rank-sum test, restricted to cell types with $n > 5$ cells in both groups. For the retinal atlas, which contains age-related macular degeneration samples rather than DR, the analysis focused on cell-type enrichment patterns rather than DR-specific differential expression. Statistical significance was defined at $P < 0.05$, with cell-type-level module score statistics reported in Supplementary Table 15. These enrichments provide orthogonal biological support for PRISMA axes but do not establish causal directionality.

Gene-set permutation tests were used to compute empirical P-values and reduce the risk of arbitrary gene-set enrichment. For each rank $r$, genes were assigned to rank $r$ only if their absolute loading satisfied $|A_{pr}| > |A_{pq}|$ for all $q \neq r$. Let $G_r$ denote the top genes from the rank-$r$ dominant pool and $\mathcal{B}$ the background gene pool defined as the intersection of genes expressed in the scRNA-seq dataset and candidate genes input to PRISMA after input filtering and LD pruning. Restricting the background to PRISMA candidate genes tests cell-type specificity rather than disease association.

For each of $N = 1,000$ permutations, a pseudo-gene set $G_r^{(i)}$ of size $|G_r|$ was sampled from $\mathcal{B}$ without replacement. Module scores were computed using `scanpy.tl.score_genes` with `ctrl_size=50` and `use_raw=True`, using independent random draws across iterations. For each rank $r$ and cell type $c$, the mean score difference between dataset-defined contrast groups was:

$$\Delta_{c,r}^{(i)} = \bar{S}_{c,r}^{disease} - \bar{S}_{c,r}^{control}$$

where disease/control notation refers to dataset-defined contrast labels and should not be interpreted as a healthy-control comparison for the FVM dataset. The empirical P-value was:

$$P_{empirical}(c,r) = \begin{cases} \max\left(1/N, \frac{\sum_{i=1}^{N} \mathbf{1}\left(\Delta_{c,r}^{(i)} \geq \Delta_{c,r}^{true}\right)}{N}\right), & \Delta_{c,r}^{true} \geq 0 \\ \max\left(1/N, \frac{\sum_{i=1}^{N} \mathbf{1}\left(\Delta_{c,r}^{(i)} \leq \Delta_{c,r}^{true}\right)}{N}\right), & \Delta_{c,r}^{true} < 0 \end{cases}$$

where $\Delta_{c,r}^{true}$ is the observed mean difference for the PRISMA gene set, $\mathbf{1}(\cdot)$ is the indicator function, and $1/N$ prevents P-values from reaching zero. Cell types with fewer than 10 cells in either condition were excluded. Across the three datasets and three genetic axes, nominal empirical P-values were reported and findings were considered supported when $P_{empirical} < 0.05$, the enrichment effect size exceeded the 95th percentile of the permutation null distribution, and the pattern was supported by independent datasets or sensitivity analyses.

Gene-set-size sensitivity analysis used Axis 1 dominant genes, defined by $|A_{p,0}| > |A_{p,1}|$ and $|A_{p,0}| > |A_{p,2}|$. Five cutoffs were evaluated: top 50, 100, 150, 200, and 243 genes. For each cutoff, module scores were recomputed and 1,000 permutation iterations sampled gene sets of equal size from the full transcriptome. Empirical P-values were computed as $(1 + \text{number of permutations with } \Delta \geq \text{observed})/(1 + \text{total permutations})$, with cutoff-level results reported in Supplementary Table 16.

#### Supplementary Method 8. Baseline comparison with conventional dimensionality reduction

PRISMA was benchmarked against PCA, NMF, and K-means using the same PRISMA integration-score matrix containing 8,201 SNPs across 5 tissues. All methods were configured to extract 3 components, matching the selected PRISMA rank. PCA was run with standardization using scikit-learn 1.3.0 [52]. NMF was run using scikit-learn 1.3.0 [41,52] with NNDSVD initialization, a maximum of 500 iterations, and non-negative input obtained by subtracting the global minimum and adding $1 \times 10^{-6}$. K-means was run using the Lloyd algorithm with 50 random initializations in scikit-learn 1.3.0 [52].

Tissue specificity was quantified using the Gini coefficient. For a component with tissue weights $w_1, w_2, \ldots, w_T$ sorted in ascending order, the Gini coefficient is:

$$\text{Gini} = \frac{2\sum_{i=1}^{T} i \cdot w_i}{T\sum_{i=1}^{T} w_i} - \frac{T+1}{T}$$

A Gini coefficient of 0 indicates uniform distribution across tissues, whereas a value of 1 indicates complete dominance by one tissue. Mean Gini coefficients were computed across components for each method.

Biological enrichment benchmarking compared PRISMA-derived gene sets against baseline-derived gene sets using independent scRNA-seq datasets. For PRISMA, the top 200 genes ranked by absolute factor loading per component were used for module scoring in FVM (GSE165784) [46] and PBMC (GSE248284) [47]. Module scores were computed using `scanpy.tl.score_genes` (scanpy 1.9.3) [49] with `ctrl_size=50` and `use_raw=True`. Significance was assessed by permutation testing with 1,000 iterations, using empirical P-values defined as the fraction of random gene sets producing equal or stronger enrichment than the observed gene set.

#### Supplementary Method 9. Height GWAS negative-control analysis

To evaluate trait-dependent regulatory reprioritization, the identical PRISMA pipeline was applied to adult height GWAS summary statistics from GWAS Catalog accession ebi-a-GCST90018959 [12,13], corresponding to European ancestry and N=360,388 [12]. The same five-tissue eQTL panel was used: retina, artery, blood, pancreas, and brain. Height was selected as a negative control because it is a highly polygenic anthropometric trait with genetic architecture distinct from diabetic retinopathy.

The height tensor was decomposed using PRISMA with R=3, matching the DR analysis. Decomposed ranks were matched by dominant tissue loading, for example by aligning the DR vascular-metabolic axis with the height tissue-matched axis. The top 200 genes from each height axis were then compared against the corresponding DR axis. Gene-set similarity was quantified by Jaccard index:

$$J(A, B) = \frac{|A \cap B|}{|A \cup B|}$$

where $A$ and $B$ represent the top 200 gene sets from height and DR. Top-200 overlap fractions quantify shared gene membership, while Jaccard indices account for the union of the two gene sets. Hypergeometric tests were used to assess whether observed overlaps exceeded expectation relative to the shared eQTL-mappable gene universe. Trait-dependent rank-shift analysis then examined whether high-weight genes in one trait retained their prioritization in the other. Representative rank-shift records are reported in Supplementary Table 19. Tissue specificity, using the Gini coefficient, and CORCONDIA were also computed for the height decomposition.

#### Supplementary Method 10. Exploratory vitreous proteomics and metabolomics

Exploratory vitreous multi-omics analyses compared PRISMA factor scores with matched proteomic and metabolomic profiles from vitreous humor specimens. The cohort comprised 13 vitrectomy samples: 7 severe diabetic retinopathy samples and 6 non-diabetic controls. Targeted proteomics quantified 429 proteins whose encoding genes overlapped the union of PRISMA factor-loading gene sets across all three trajectories.

For proteomic module scoring, each protein was compared between DR and control samples using a two-sided Mann-Whitney U test. Proteins with $P < 0.05$ were mapped back to the PRISMA trajectory from which their encoding gene was drawn, yielding trajectory-specific significant-protein sets. Per-sample module scores were calculated as the mean row-wise Z-score of significant proteins within each trajectory's gene set, with Z-score standardization performed across all 13 samples for each protein. Module-level differences between DR and control groups were tested using a two-sided Mann-Whitney U test, with module-score and target-level proteomic statistics reported in Supplementary Table 17.

Untargeted metabolomics profiling of the same specimens produced annotated metabolomic features after annotation-level filtering of likely exogenous compounds, including drugs, pesticides, and industrial chemicals, using a keyword-based filter. Because untargeted annotations remain putative, individual compound names were interpreted as annotated features rather than definitive molecular identities. To isolate within-disease heterogeneity, metabolite-axis correlation analysis was restricted to the 7 DR samples. For each retained feature, Spearman rank correlations were computed against each of the three trajectory module scores.

Multiple testing was controlled by Benjamini-Hochberg FDR correction [50] independently for each trajectory axis. For each axis, P-values from Spearman correlations across metabolites were adjusted using `statsmodels.stats.multitest.multipletests()` in Python 3.10 [51]. Metabolites with FDR $q < 0.05$ were considered statistically significant, and those with $0.05 \leq q < 0.10$ were retained as exploratory supplementary findings. Given the small DR metabolomics sample size (n=7), effect size and sensitivity-filtering behavior were prioritized alongside nominal statistical significance.

These correlations were used to nominate candidate molecular correlates of PRISMA axes. They do not establish causality, temporal ordering, stable biomarkers, or clinical utility.

For each metabolite, the trajectory with the largest absolute correlation was designated as the primary driver. Trajectory specificity was quantified as:

$$S_m = \max_r |\rho_{mr}| - \mathbf{secondmax}_r |\rho_{mr}|$$

where $\rho_{mr}$ is the Spearman correlation between metabolite $m$ and trajectory $r$. This score was defined for metabolites where $\max_r |\rho_{mr}| \neq 0$; metabolites with $|\rho_{max}| < 0.75$ were excluded from specificity analysis. A high $S_m$ indicates preferential association with one trajectory, while a low $S_m$ indicates distributed association across trajectories, consistent with shared downstream biological convergence.

Leave-one-out cross-validation (LOOCV) was performed across the 7 DR samples. For each held-out patient, the Spearman correlation between a metabolite and its primary-driver trajectory score was recomputed on the remaining 6 patients. Metabolites were retained for final exploratory analysis only if they satisfied: FDR $q < 0.10$ for the primary axis, primary-driver $|\rho| > 0.75$, LOOCV mean correlation > 0.5, and LOOCV standard deviation < 0.25. This multi-criteria filter was used as an exploratory sensitivity screen to prioritize candidate metabolite-axis associations for future evaluation.

### Supplementary Method 11. Post-GWAS comparator analyses

#### 11.1 Comparator target definition

Post-GWAS comparator analyses were performed to evaluate gene-level concordance between PRISMA-prioritized targets and established GWAS gene-prioritization frameworks. These analyses were not used to fit PRISMA and were not interpreted as causal validation of all PRISMA targets.

Supplementary Table 5 contains 549 PRISMA target rows. Because established comparator methods such as MAGMA, SMR/HEIDI, and coloc operate at gene or gene-tissue levels rather than duplicated target-row levels, the 549 PRISMA target rows were projected to 405 unique gene symbols before comparator analysis. This 405-gene projection was used as the primary PRISMA comparator target set. Axis-specific comparator gene sets were constructed from the same unique-gene projection by assigning genes to their primary PRISMA axis: vascular-metabolic, circulating immune-inflammatory, or retina-specific neurodegenerative. An exploratory subthreshold set was also defined from PRISMA-prioritized genes that did not pass the conventional genome-wide significance threshold. This subthreshold set was used only for sensitivity analyses and did not define primary causal claims.

#### 11.2 MAGMA analysis

MAGMA was used as a GWAS-only comparator to assess gene-level and gene-set burden among PRISMA-prioritized targets. Diabetic retinopathy GWAS meta-analysis summary statistics were formatted for MAGMA using SNP identifiers, chromosome, base-pair position, P values, and sample size. Analyses used GRCh37/hg19 coordinates, a 1000 Genomes European PLINK LD reference, MAGMA v1.10, and the NCBI37.3 gene-location file. SNP-to-gene annotation was generated with the MAGMA `--annotate` step using the prepared SNP-location file and NCBI37.3 gene locations. Gene-based analysis was then performed using the 1000 Genomes European LD reference and prepared GWAS P-value file, followed by competitive gene-set analysis for the PRISMA target set, the three axis-specific gene sets, and the exploratory subthreshold set.

MAGMA gene-set testing used Entrez-ID GMT files because MAGMA gene results are indexed by Entrez identifiers from the NCBI37.3 gene-location file. The original readable gene-symbol GMT was therefore mapped to Entrez identifiers before the final MAGMA gene-set run. Genes not mappable to MAGMA Entrez identifiers were excluded from MAGMA gene-set testing and reported as a mapping limitation. Specifically, 25 of 405 PRISMA unique genes did not map to the NCBI37.3 MAGMA gene-location symbols. Final MAGMA gene-set results were corrected across the tested PRISMA-derived gene sets using Benjamini-Hochberg FDR correction. The PRISMA top-target gene set and all three axis-specific sets were evaluated as comparator gene sets; significant MAGMA burden was interpreted as GWAS gene-set concordance, not as evidence that MAGMA validates PRISMA axes or establishes gene causality.

##### 11.3 SMR/HEIDI comparator analysis

SMR/HEIDI was used as a formal GWAS-eQTL gene-level comparator. The primary SMR comparator used legacy all-tissue SMR outputs generated from the same diabetic retinopathy GWAS meta-analysis. Parsed tissue/source labels included Artery_Aorta, Artery_Coronary, Artery_Tibial, Brain_Cerebellum, Brain_Cortex, Kidney_Cortex, Nerve_Tibial, Pancreas, Retina, Whole_Blood, and eQTLGen. These outputs were reused as existing comparator results rather than rerun as part of PRISMA model fitting.

Strict SMR support was defined as all-tissue Benjamini-Hochberg adjusted SMR $q < 0.05$ together with HEIDI $P > 0.01$, when HEIDI was available. This all-tissue correction was applied across parsed SMR tissue/source outputs. Gene-level enrichment of strict SMR-supported genes among PRISMA-prioritized genes was evaluated within the PRISMA background universe of 7,507 genes. Among strict all-tissue SMR-supported genes, 123 were within the PRISMA background and 31 overlapped the 405-gene PRISMA comparator target set. Overlap enrichment was evaluated using a background-restricted hypergeometric test and an approximate odds ratio from the corresponding 2 x 2 contingency table.

The legacy Retina SMR output used an external institutional retina eQTL source and was not the same retina eQTL input used for the manuscript PRISMA decomposition. Therefore, retina-source caveats were tracked explicitly. Manuscript-aligned Retina SMR, when retained, was handled as a separate sensitivity analysis rather than merged unqualified with the legacy all-tissue SMR comparator. SMR axis-tissue consistency was also treated as exploratory because the all-tissue SMR comparator was designed primarily to assess gene-level concordance rather than to validate PRISMA tissue axes.

##### 11.4 Targeted colocalization analysis

Targeted colocalization was performed as a shared-variant sensitivity analysis conditioned on strict SMR-PRISMA overlap. Colocalization was not performed genome-wide and was not used as an unbiased scan for all PRISMA targets. The target set consisted of 31 genes that were both in the PRISMA 405-gene comparator projection and supported by strict all-tissue SMR/HEIDI. Candidate coloc tests were then defined at the gene-tissue-cis-window level.

Stage 1 coloc input QC evaluated each strict SMR-PRISMA gene-tissue pair. A pair was considered runnable only if a local eQTL summary was available for the exact tissue, eQTL sample size was available, GWAS sample size and diabetic retinopathy case fraction were documented, at least 50 shared SNPs remained after harmonization, and allele harmonization passed QC. Exact local eQTL summaries were available for Artery_Tibial, Brain_Cortex, Pancreas, and Whole_Blood. Tissues without exact local summaries were not substituted with related tissues. Retina was treated as sensitivity-only because its coloc-ready sample-size metadata were not available in the formal GTEx-based coloc workflow.

For each gene-tissue pair, coloc input windows were defined as probe base-pair position $\pm 500$ kb. When the probe coordinate was missing, the top SMR SNP coordinate was used as a fallback and flagged. SNP matching used rsID where available, with chromosome-position matching used only as a fallback. GWAS and eQTL alleles were harmonized before coloc input generation. eQTL beta values were sign-flipped when alleles were reversed, strand-ambiguous or unharmonizable SNPs were removed, and variance terms were computed as squared standard errors. Minor allele frequency was derived from harmonized GWAS effect-allele frequency as $\min(EAF, 1 - EAF)$ unless eQTL MAF was available; this source was recorded as a GWAS EAF proxy. The diabetic retinopathy GWAS was analyzed as a case-control dataset using the documented meta-analysis case fraction.

After Stage 1 QC, 23 gene-tissue pairs were classified as runnable. Pairs were excluded from the formal targeted coloc run if they lacked exact local eQTL summaries, had fewer than 50 shared harmonized SNPs, or had ambiguous allele harmonization. Stage 2 targeted coloc used `coloc.abf` only for these 23 QC-passed pairs. Colocalization support was defined as PP.H4 $\geq 0.80$; 0.50 $\leq$ PP.H4 < 0.80 was treated as suggestive; PP.H4 < 0.50 was interpreted as no coloc support in that tested tissue-window. These thresholds were used for sensitivity interpretation only. Targeted coloc was interpreted as local shared-variant support for selected strict SMR-supported PRISMA genes, not as causal proof, not as validation of all targets, and not as evidence that PRISMA replaces formal fine-mapping or colocalization analyses.

#### 11.5 Mixed sensitivity analyses

Two additional sensitivity analyses were considered exploratory and were not used as primary main-text claims. First, SMR axis-tissue consistency was evaluated by projecting strict SMR-supported PRISMA genes back onto PRISMA axes and comparing supported tissues with broad axis expectations. These results were mixed and were used only to contextualize gene-level SMR concordance. They were not interpreted as validation of PRISMA axes.

Second, subthreshold PRISMA targets were evaluated in single-cell enrichment sensitivity analyses. Subthreshold analyses used the 405-gene projection and PRISMA background universe to construct pooled and axis-specific subthreshold gene sets. Module scores and permutation tests were run against the same single-cell resources used for the main orthogonal validation analyses. The subthreshold analysis was retained as supplementary because enrichment was mixed: the Axis 1 subthreshold set showed FVM fibroblast enrichment, whereas Axis 2 PBMC and Axis 3 retinal compartment results were not consistently significant. This analysis supports exploratory biological coherence for a subset of subthreshold PRISMA signal but does not establish causality or validate every subthreshold target.
