## supplementary notes for "PRISMA: A tensor-based framework for deconstructing the genetic architecture of complex diseases, with application to diabetic retinopathy": Supplementary_Methods.md

# Supplementary Methods

## Metabolomics compound filtering

Untargeted vitreous metabolomics features were filtered before trajectory-correlation analysis to prioritize endogenous metabolites. Compounds were excluded if their names or annotations indicated likely exogenous origin, including therapeutic drugs, pesticides, herbicides, industrial chemicals, polymer additives, food additives, cosmetic ingredients, and laboratory contaminants. Remaining metabolites were retained for downstream Spearman correlation analysis against PRISMA trajectory module scores in the diabetic retinopathy samples.

The filtering step was applied before multiple-testing correction. For each PRISMA axis, raw Spearman correlation P-values across retained metabolites were adjusted using the Benjamini-Hochberg procedure. Robust metabolite correlates were then selected using the criteria described in the main Methods: primary-axis |rho| > 0.75, leave-one-out cross-validation mean correlation > 0.5, leave-one-out standard deviation < 0.25, and FDR q < 0.10 for exploratory reporting.
