## supplementary notes for "PRISMA: A tensor-based framework for deconstructing the genetic architecture of complex diseases, with application to diabetic retinopathy": Supplementary_Note_1_Simulation_Validation.md

# Supplementary Note 1: Monte Carlo Simulation Validation

## Overview

To validate PRISMA's capacity to systematically disentangle overlapping pleiotropic effects, we conducted extensive Monte Carlo benchmarking simulations using a synthetic high-dimensional variant-by-tissue association array.

## Simulation Design

We engineered a ground-truth multi-trajectory architecture consisting of two distinct latent pathogenic axes. Each axis was strictly driven by an independent set of genetic variants operating within specific tissue profiles. Crucially, to mirror the true input of our framework, the simulated data were generated using the exact Summary-data-based Mendelian Randomization (SMR) statistical model, coupling underlying GWAS and eQTL effects. 

To rigorously evaluate the model's specificity and robustness against overfitting, we intentionally embedded a high proportion of null variants (representing pure background noise) and applied substantial global Gaussian noise to the entire array.

## Results

### Trajectory Recovery

Despite this high-noise environment, a representative realization demonstrated that PRISMA robustly recovered the latent multi-trajectory architecture. As shown in the simulation outputs supporting Fig. 2, the algorithm accurately delineated the distinct tissue specificities corresponding to the ground truth. It achieved a clean, block-diagonal separation of tissue weights across the two axes, confirming that PRISMA retains high spatial resolution without introducing signal cross-talk between distinct tissue modules. 

Furthermore, the locus-level factor loadings (Factor A) precisely captured the specific genetic variants driving each respective axis, effectively suppressing the signals from the embedded null variants to near zero.

### Type I Error Calibration

To systematically quantify this statistical rigor, we performed hundreds of Monte Carlo iterations. We first evaluated the empirical false positive rate (Type I error) against an independently derived global null distribution using completely null datasets. PRISMA maintained conservative calibration across standard nominal thresholds (e.g., $\alpha = 0.05$), demonstrating that the algorithm does not inflate signals or artificially hallucinate latent structure from pure background noise (Fig. 2a).

### Statistical Power

Moreover, strict SNP-level power curve analysis across varying signal-to-noise ratios revealed a highly responsive sigmoidal trajectory (Fig. 2b). In the frozen April 18 simulation benchmark, PRISMA approached the 80% power threshold at signal level 2.5 (78.96%) and exceeded it at signal level 3.5 (95.25%), quickly asymptoting to complete recovery at stronger signal levels.

## Conclusion

Together, these quantitative benchmarking results establish that PRISMA's advanced array decomposition robustly isolates true biological trajectories from complex genetic noise, providing an unshakeable statistical foundation for its application to real-world clinical data.

---

**Fig. 2.** Monte Carlo simulation validation of PRISMA. The main figure summarizes Type I error calibration, statistical power, SNP recovery precision, and computational efficiency.
