## supplementary notes for "PRISMA: A tensor-based framework for deconstructing the genetic architecture of complex diseases, with application to diabetic retinopathy": Supplementary_Note_2_Methodological_Comparisons.md

# Supplementary Note 2: Methodological Comparisons and Theoretical Properties

## Positioning PRISMA in the Methods Landscape

PRISMA complements both aggregate tissue-level heritability partitioning 
(e.g., stratified LD score regression, S-LDSC) and variant-level 
fine-mapping (e.g., mvSuSiE) by providing trajectory-resolved decomposition 
at the locus-by-tissue level. While S-LDSC identifies which tissues are 
enriched for disease heritability in aggregate, and while mvSuSiE identifies 
which independent variants causally drive pleiotropic traits with high 
resolution, PRISMA explicitly resolves how these loci orchestrate disease 
heterogeneity through specific cross-tissue regulatory networks, 
deconvolving aggregate tissue-level enrichment into mechanistically distinct 
pathogenic axes. For example, where S-LDSC might reveal that "whole blood" 
is enriched for DR heritability, PRISMA further resolves this into two 
independent trajectories---a vascular-metabolic axis (Axis 1) and an 
immune-inflammatory axis (Axis 2)---each with distinct driving variants, 
cellular mechanisms, and patient-level genetic burden scores.

## Mathematical Advantages of the PRISMA Framework

A core mathematical advantage of our framework lies in the integration of 
non-negativity constraints with graph Laplacian regularization within the 
CP decomposition. The non-negativity constraints ensure biologically 
interpretable factor loadings (all weights are positive contributions), 
while the graph Laplacian preserves local linkage disequilibrium structure 
and encourages SNPs in high LD to share similar loadings. This design 
naturally accommodates biological pathways that converge---as mirrored by 
the DR funnel model---allowing PRISMA to capture genetically distinct 
upstream axes that may converge on shared terminal pathogenic routes.

## Graph Laplacian vs. Standard Factorization

A potential concern is whether PRISMA's tissue-specificity advantage stems primarily from the graph Laplacian regularization rather than the coupled block-wise architecture. While graph-regularized NMF has been applied to single-matrix problems, PRISMA's innovation lies in the **coupled decomposition across genomic blocks with shared tissue factors (B)**, which enables borrowing of information across loci while preserving local LD structure. The graph Laplacian ensures that SNPs in high LD share coordinated loadings within each block, while the shared B matrix enforces consistent tissue-specificity patterns across the genome. This dual-level structure—local LD preservation combined with global tissue coherence—is not achievable by applying graph regularization to a single concatenated matrix, as the latter would lose the block-wise genomic topology that is fundamental to capturing locus-specific regulatory effects.

## Joint Multi-Tissue Decomposition

Furthermore, by decomposing the multi-tissue association tensor jointly, 
PRISMA captures coordinated cross-tissue regulatory patterns as coherent 
latent factors, rather than treating each tissue as an independent 
analytical unit. This joint decomposition enables the framework to identify 
tissue-specific trajectories even when individual tissues have limited 
sample sizes, by borrowing strength across the shared genomic architecture.

## Comparison with Multi-Omics Integration Methods

### MOFA+ (Multi-Omics Factor Analysis)

MOFA+ performs unsupervised factor analysis to integrate proteomics, metabolomics, and transcriptomics data, discovering latent factors that capture coordinated variation across molecular layers. While MOFA+ demonstrates the power of latent factor models for biological data integration, it faces practical limitations in the context of complex disease genetics:

1. **Data requirements**: MOFA+ typically requires measured multi-omics data from the exact same individuals, which are scarce and rarely reach the massive sample sizes of modern GWAS (N>500,000).

2. **Disease association**: MOFA+ decomposes multi-modal molecular data without directly incorporating disease associations from GWAS.

3. **LD structure**: MOFA+ does not incorporate the block-wise linkage disequilibrium (LD) structure inherent to genetic data.

In contrast, PRISMA operates entirely on publicly available summary-level data (GWAS + eQTL), explicitly integrates disease associations, and preserves local LD topology through graph Laplacian regularization.

### sn-spMF (Sparse Non-negative Matrix Factorization)

sn-spMF applies sparse matrix factorization to GTEx data to learn patterns of tissue-sharing in eQTL effects. While this approach identifies tissue-specific eQTL patterns, it has key differences from PRISMA:

1. **Disease focus**: sn-spMF decomposes eQTL effect matrices without directly incorporating disease associations, whereas PRISMA integrates GWAS signals to identify disease-relevant trajectories.

2. **LD preservation**: sn-spMF treats variants as independent observations, whereas PRISMA explicitly preserves local LD structure through graph Laplacian regularization.

3. **Interpretability**: sn-spMF produces tissue-sharing patterns, whereas PRISMA produces patient-level genetic burden scores for each trajectory, enabling direct clinical translation.

## Theoretical Properties

### Convergence Guarantees

The block coordinate descent framework with convex subproblems has established convergence guarantees under mild regularity conditions (Bertsekas, 1999; Tseng, 2001). While we do not provide a formal convergence proof for the combined Graph Laplacian and non-negativity constraints, empirical validation across 50 random initializations demonstrated highly stable solutions (mean pairwise cosine similarity = 1.000 ± 7×10⁻¹⁷ for Factor A, 1.000 ± 2×10⁻¹⁷ for Factor B, 1.000 ± 0 for Factor C), confirming that the LD-informed regularization effectively constrains the solution space.

### Identifiability

For standard CP decomposition, the Kruskal condition guarantees unique recovery of latent factors. However, in our single-phenotype setting where Factor C serves as a scaling vector, the model reduces to coupled matrix factorization, and the Kruskal condition degenerates. We ensure solution uniqueness and stability through three complementary mechanisms:

1. **Non-negativity constraints**: Restricting all factor loadings to non-negative values constrains the solution space to biologically interpretable positive contributions, eliminating sign ambiguity.

2. **Graph Laplacian regularization**: The LD-informed penalty enforces local genetic structure, preventing arbitrary solutions by requiring SNPs in high LD to share coordinated loadings.

3. **Empirical stability**: Across 50 random initializations, solutions converged to nearly identical factor matrices (cosine similarity ≈ 1.000), demonstrating that the biological priors effectively stabilize the decomposition.

## Computational Efficiency

PRISMA's block-wise alternating least squares (ALS) architecture enables efficient processing of genome-wide data. In five repeated full real-data runs, the framework processed 993,226 pre-LD candidate variants across five eQTL tissues (4,966,130 variant-by-tissue cells) with a median wall-clock runtime of 206.3 seconds on a standard consumer-grade processor (Intel Core i9 14900K; Supplementary Table 22), demonstrating scalability to biobank-scale datasets without requiring high-performance computing infrastructure.

## References

Bertsekas, D. P. (1999). *Nonlinear Programming* (2nd ed.). Athena Scientific.

Tseng, P. (2001). Convergence of a block coordinate descent method for nondifferentiable minimization. *Journal of Optimization Theory and Applications*, 109(3), 475-494.
