## supplementary table legends for "PRISMA: A tensor-based framework for deconstructing the genetic architecture of complex diseases, with application to diabetic retinopathy"

# Supplementary Table Legends

## Supplementary Table 1 | GWAS summary statistics metadata

Metadata for genome-wide association study (GWAS) datasets used in PRISMA tensor decomposition. Includes sample sizes, ancestry, phenotype definitions, and data sources for diabetic retinopathy (DR) GWAS from FinnGen and Million Veteran Program (MVP).

---

## Supplementary Table 2 | Multi-tissue eQTL metadata

Metadata for expression quantitative trait loci (eQTL) datasets across five DR-relevant tissues: retina, tibial artery, brain cortex, pancreas, and whole blood. Includes sample sizes, tissue sources, and data provenance from the human retina eQTL atlas and SMR-formatted GTEx v8 BESD files. GTEx tissue rows refer to BESD-derived SNP-level representative eQTL inputs used for the manuscript analysis, not raw GTEx Portal allpairs tables.

---

## Supplementary Table 3 | Single-cell RNA-seq metadata

Metadata for three independent single-cell transcriptomic datasets used for PRISMA validation: fibrovascular membrane (FVM, GSE165784), peripheral blood mononuclear cells (PBMC, GSE248284), and human retina (GSE137537). Includes cell counts, disease status, and data sources.

---

## Supplementary Table 4 | DR top genes across three axes

Top 200 genes per axis ranked by absolute PRISMA factor loadings (Factor A). Includes gene identifiers, weights, sentinel SNP labels, and within-axis ranks for the three genetic trajectories: Axis 0 (vascular-metabolic), Axis 1 (immune-inflammatory), and Axis 2 (retinal-neural).

---

## Supplementary Table 5 | Complete 549 PRISMA prioritized targets

Complete list of 549 genes prioritized by PRISMA across three genetic axes, representing the top 200 genes per axis that successfully mapped to genomic positions and GWAS signals. This table resolves the apparent discrepancy between "top 200 per axis" (600 total) and "549 targets": the difference arises from genes lacking genomic position annotations or GWAS coverage.

Columns include gene symbol, assigned axis, PRISMA factor loading, chromosome, hg19 genomic position, sub-threshold status relative to the conventional genome-wide significance threshold, and rank within axis.

Summary: Axis 0 (vascular-metabolic), 184 genes; Axis 1 (immune-inflammatory), 189 genes; Axis 2 (retinal-neural), 176 genes; total, 549 genes; sub-threshold, 403 genes (73%). This table provides the gene-level resolution underlying Figure 4 and supports the manuscript's claim that PRISMA prioritizes sub-threshold polygenic signals through tissue-specific regulatory networks.

---

## Supplementary Table 6 | Height GWAS top genes (negative control)

Top 200 genes per axis from height GWAS analysis (GWAS Catalog accession ebi-a-GCST90018959, corresponding to the European-ancestry adult height GWAS from Sakaue et al. (N=360,388)) using the identical PRISMA framework. Demonstrates trait-specific gene prioritization despite a shared tissue-regulatory backbone.

---

## Supplementary Table 7 | Baseline method comparison

Quantitative comparison of PRISMA against conventional dimensionality reduction methods (PCA, NMF, K-means). Includes Gini coefficients for tissue specificity and Factor B tissue weight matrices demonstrating PRISMA's superior tissue-regulatory resolution. The `Record_Type` column distinguishes method-level Gini summaries from tissue-weight matrix rows. PRISMA rank order follows the current manuscript and Supplementary Table 12: Rank 0 = Axis 1 (vascular-metabolic), Rank 1 = Axis 2 (immune-inflammatory), and Rank 2 = Axis 3 (retina-specific neurodegenerative).

---

## Supplementary Table 8 | Permutation test results

Cell-type-specific enrichment statistics from direction-aware gene-set permutation tests (1,000 iterations) across FVM and PBMC datasets for all three PRISMA axes. Includes observed mean differences, signed enrichment scores, permutation null summaries, empirical P-values, and direction flags. Positive `True_Mean_Diff` values indicate enrichment in disease cells, whereas negative values indicate depletion-direction effects.

---

## Supplementary Table 9 | Vitreous metabolomics robust DR-only correlates

Axis-associated metabolomic features in vitreous humor from severe proliferative DR patients (n=7), filtered for robust exploratory association. Includes metabolite class annotations, assigned PRISMA axis, primary-axis Spearman rho, primary-axis FDR q-value, orthogonal specificity score, leave-one-out cross-validation metrics, and axis-wise rho/q values. The generally low specificity scores across prioritized features are reported explicitly because they support the downstream-convergence model: genetically and spatially distinct PRISMA axes converge onto a shared vitreous metabolic milieu, with Axis 1 (vascular-metabolic) accounting for most robust macroscopic metabolic variance in this fluid compartment.

---

## Supplementary Table 10 | Lambda sensitivity analysis

Adjusted Rand Index (ARI) of axis-to-tissue assignments across 10,000-fold variation in the graph Laplacian penalty weight (lambda in {0.01, 0.1, 1, 10, 100}). Demonstrates perfect stability (ARI = 1.00) of PRISMA decomposition across extreme hyperparameter perturbations.

---

## Supplementary Table 11 | Vitreous metabolomics full FDR results

Complete metabolomics dataset with FDR-corrected correlations for all retained metabolites across three PRISMA axes. Includes compound identifiers, chemical classes, Spearman correlations, raw P-values, and FDR q-values for each axis.

---

## Supplementary Table 12 | PRISMA Factor B and Factor C matrices

Global factor matrices from the final PRISMA decomposition. Factor B reports tissue-mode loadings for each latent axis across retina, artery, cortex, pancreas, and whole blood. Factor C reports the phenotype-mode scaling vector for the current single-phenotype diabetic retinopathy application. These matrices support the axis-level tissue interpretation in the Results and document the single-phenotype scaling used in the coupled matrix factorization regime.

---

## Supplementary Table 13 | Rank selection and model fit across candidate ranks

Model fit statistics for PRISMA decompositions evaluated across ranks R = 1 to 5. Includes variance explained and Core Consistency Diagnostic (CORCONDIA) values used to select the final rank-3 decomposition.

---

## Supplementary Table 14 | Convergence stability across 50 random initializations

Summary of pairwise cosine similarities for Factor A, Factor B, and Factor C across 50 independent random initializations after optimal component alignment. Reports mean, standard deviation, minimum, and maximum pairwise cosine similarity for each factor matrix.

---

## Supplementary Table 15 | Single-cell module score statistics across validation atlases

Cell-type-level PRISMA module score statistics across fibrovascular membrane (FVM), peripheral blood mononuclear cell (PBMC), and retinal single-cell atlases. FVM and PBMC rows report dataset-defined contrast-group mean differences. FVM contrasts should be interpreted as within-fibrovascular-membrane cell-type/module-score contrasts rather than healthy-control comparisons, whereas PBMC rows report DR-versus-control contrasts where available. Retina rows report rank-2 cell-type enrichment scores, empirical P-values, and cell counts, including the neuron-versus-non-neuron validation summary.

---

## Supplementary Table 16 | Gene-set size sensitivity analysis for Axis 1 FVM enrichment

Permutation-based sensitivity analysis testing whether Axis 1 enrichment in FVM pathological cells is robust to gene-set size. Reports empirical P-values across sliding cutoffs from the top 50 to all 243 strictly Axis 1-dominant targets.

---

## Supplementary Table 17 | Vitreous proteomics module scores and target-level differential statistics

Proteomics validation table for the vitreous humor cohort. Module-score rows report per-sample PRISMA proteomic module Z-scores across Rank 0, Rank 1, and Rank 2. Module-differential rows report exact two-sided Mann-Whitney U tests comparing DR versus control module scores. Target-statistic rows report DR-versus-control differential abundance statistics for the 429 PRISMA-overlapping vitreous proteins, with additional records for top PRISMA targets where factor weights are available.

---

## Supplementary Table 18 | Simulation benchmark results

Structured benchmark records underlying the PRISMA simulation validation. Includes Type I error calibration, statistical power across effect sizes, ROC-derived SNP recovery performance, and runtime scaling values. These records use the frozen April 18 enhanced Figure 2 code path and are linked to the full Figure 2 source data in Supplementary Data 3.

---

## Supplementary Table 19 | Height negative-control comparison statistics

Negative-control comparison between diabetic retinopathy and adult height PRISMA decompositions. Includes model-fit summaries, axis-wise tissue-weight summaries, top-gene overlap and hypergeometric enrichment statistics, gene-symbol annotations for set-membership records, and representative gene-level rank-shift records supporting the trait-specific reprioritization analysis. Positive `Rank_Shift_Height_minus_DR` values indicate stronger DR prioritization, whereas negative values indicate stronger Height prioritization.

---

## Supplementary Table 20 | PRISMA weight-biological effect correlation analysis

Gene-level records supporting the correlation between PRISMA mathematical weights and in vivo disease dysregulation in fibrovascular membrane structural cells. Includes PRISMA axis assignments, factor weights, differential-expression effect sizes, and source annotations for the weight-biology comparison shown in Figure 5a.

---

## Supplementary Table 21 | Lambda sensitivity factor-weight trajectories

Detailed factor-weight trajectories across graph Laplacian penalty values. Reports raw rank identities, aligned biological axes, tissue-mode weights, dominant tissue weights, and ARI summaries, enabling direct verification that tissue-axis identities and dominant weights remain stable across the tested lambda range.

---

## Supplementary Table 22 | Real-data computational runtime benchmark

Five repeated full real-data PRISMA runs using the manuscript input manifest, rank 3, lambda = 0.1, and 20 ALS iterations. The table reports raw backbone variants, pre-LD candidate variants, pre-LD variant-by-tissue cells, final gene-representative variants, LD blocks, wall-clock runtime, CPU time, peak memory, software environment, and the exact command used for each run.

---

## Supplementary Table 23 | Expanded summary of PRISMA-derived diabetic retinopathy axes

Expanded editable version of the concise main-text Table 1. This table summarizes the three PRISMA-derived diabetic retinopathy axes, including trajectory labels, dominant tissue loadings, regulatory interpretations, orthogonal single-cell support, exploratory vitreous molecular patterns, scope-of-inference boundaries, and links to supporting supplementary resources. The table is intended as a reader-facing summary and does not introduce new analyses or numerical results beyond those reported in the main text and supporting supplementary tables.

## Supplementary Table 24 | Post-GWAS comparator summary

Summary of post-GWAS comparator analyses added after PRISMA decomposition, including target projection, MAGMA gene-set burden, SMR/HEIDI gene-level concordance, targeted coloc shared-variant sensitivity, and exploratory subthreshold scRNA sensitivity. Comparator analyses use the 405 unique-gene projection of the 549 PRISMA target rows where gene-level methods require unique gene symbols.

## Supplementary Table 25 | MAGMA post-GWAS results

MAGMA gene-set burden and gene-level overlap results for the PRISMA 405-gene projection, axis-specific gene sets, and exploratory subthreshold gene set. MAGMA analyses used GRCh37 coordinates, 1000 Genomes European LD reference data, MAGMA v1.10, NCBI37.3 gene locations, and Entrez-ID gene-set files. Genes not mappable to MAGMA Entrez identifiers were excluded from MAGMA gene-set testing and reported as a mapping limitation.

## Supplementary Table 26 | SMR/HEIDI concordance

Strict SMR/HEIDI-supported genes overlapping the PRISMA 405-gene projection. Strict support was defined as all-tissue BH-adjusted SMR q < 0.05 and HEIDI P > 0.01. The table reports best SMR tissue, tested tissues, PRISMA axis assignment, subthreshold status, and strict or lenient tissue support. Tissue-source caveats, especially for legacy Retina SMR, are described in Supplementary Method 11.

## Supplementary Table 27 | Targeted coloc sensitivity analysis

Targeted coloc.abf results for QC-passed strict SMR-PRISMA gene-tissue pairs, together with axis and tissue summaries. Colocalization was not performed genome-wide; it was restricted to eligible gene-tissue cis-window pairs after Stage 1 QC. PP.H4 ≥ 0.80 was considered coloc-supported shared-variant evidence, 0.50 ≤ PP.H4 < 0.80 was treated as suggestive, and PP.H4 < 0.50 was interpreted as no coloc support in the tested tissue-window.

## Supplementary Table 28 | Exploratory subthreshold scRNA sensitivity

Expected-compartment summary for exploratory subthreshold PRISMA single-cell RNA-seq module-score and permutation sensitivity analyses. The table reports axis-specific subthreshold gene sets, expected datasets and cell compartments, detected gene counts, effect sizes, empirical P values, FDR-adjusted module-score statistics, and support levels. These results are supplementary sensitivity analyses and are not used as primary main-text claims.

---

**Note**: Supplementary Tables are provided as CSV or XLSX files for computational reproducibility and reader-facing review. All tables use standard column naming conventions and are compatible with R, Python, and Excel where applicable.
